# Structural Basis for Potent Neutralization of Betacoronaviruses by Single-domain Camelid Antibodies

**DOI:** 10.1101/2020.03.26.010165

**Authors:** Daniel Wrapp, Dorien De Vlieger, Kizzmekia S. Corbett, Gretel M. Torres, Wander Van Breedam, Kenny Roose, Loes van Schie, VIB-CMB COVID-19 Response Team, Markus Hoffmann, Stefan Pöhlmann, Barney S. Graham, Nico Callewaert, Bert Schepens, Xavier Saelens, Jason S. McLellan

**Author notes:** Correspondence (B.S.), (X.S.), (J.S.M.). These authors contributed equally to this work (Daniel Wrapp, Dorien De Vlieger).

## Abstract

The pathogenic Middle East respiratory syndrome coronavirus (MERS-CoV), severe acute respiratory syndrome coronavirus (SARS-CoV-1) and COVID-19 coronavirus (SARS-CoV-2) have all emerged into the human population with devastating consequences. These viruses make use of a large envelope protein called spike (S) to engage host cell receptors and catalyze membrane fusion. Because of the vital role that these S proteins play, they represent a vulnerable target for the development of therapeutics to combat these highly pathogenic coronaviruses. Here, we describe the isolation and characterization of single-domain antibodies (VHHs) from a llama immunized with prefusion-stabilized coronavirus spikes. These VHHs are capable of potently neutralizing MERS-CoV or SARS-CoV-1 S pseudotyped viruses. The crystal structures of these VHHs bound to their respective viral targets reveal two distinct epitopes, but both VHHs block receptor binding. We also show cross-reactivity between the SARS-CoV-1 S-directed VHH and SARS-CoV-2 S, and demonstrate that this cross-reactive VHH is capable of neutralizing SARS-CoV-2 S pseudotyped viruses as a bivalent human IgG Fc-fusion. These data provide a molecular basis for the neutralization of pathogenic betacoronaviruses by VHHs and suggest that these molecules may serve as useful therapeutics during coronavirus outbreaks.

## INTRODUCTION

Coronaviruses are enveloped, positive-sense RNA viruses that are divided into four genera (α, β, γ, δ) and infect a wide variety of host organisms (Woo et al., 2009). There are at least seven coronaviruses that can cause disease in humans, and four of these viruses (HCoV-HKU1, HCoV-OC43, HCoV-NL63 and HCoV-229E) circulate seasonally throughout the global population, causing mild respiratory disease in most patients (Gaunt et al., 2010). The three remaining viruses, SARS-CoV-1, MERS-CoV and SARS-CoV-2, are zoonotic pathogens that have caused epidemics or pandemics with severe and often fatal symptoms after emerging into the human population (Chan et al., 2020; Huang et al., 2020; Ksiazek et al., 2003; Lu et al., 2020; Zaki et al., 2012). For these highly pathogenic betacoronaviruses, prophylactics and therapeutic treatments are needed.

The surfaces of coronaviruses are decorated with a spike glycoprotein (S), a large class I fusion protein (Bosch et al., 2003). The S protein forms a trimeric complex that can be functionally categorized into two distinct subunits, S1 and S2, that are separated by a protease cleavage site. The S1 subunit contains the receptor-binding domain (RBD), which interacts with a proteinaceous host-cell receptor to trigger membrane fusion. The S2 subunit contains the fusion machinery, including the hydrophobic fusion peptide and the α-helical heptad repeats. The functional host cell receptors for SARS-CoV-1 and MERS-CoV are angiotensin converting enzyme 2 (ACE2) and dipeptidyl peptidase 4 (DPP4), respectively (Li et al., 2003; Raj et al., 2013). The interactions between these receptors and their respective RBDs have been thoroughly characterized, both structurally and biophysically (Li et al., 2005; Wang et al., 2013). Recently, it has been reported that SARS-CoV-2 S also makes use of ACE2 as a functional host-cell receptor and several structures of this complex have already been reported (Hoffmann, 2020; Lan, 2020; Wan et al., 2020; Yan, 2020; Zhou et al., 2020).

Recent advances in cryo-EM have allowed researchers to determine high-resolution structures of the trimeric spike protein ectodomains and understand how S functions as a macromolecular machine (Kirchdoerfer et al., 2016; Li et al., 2005; Walls et al., 2016; Wang et al., 2013). Initial cryo-EM characterization of the SARS-CoV-1 spike revealed that the RBDs adopted at least two distinct conformations. In the “up” conformation, the RBDs could be observed jutting out away from the rest of S, such that they could easily engage ACE2 without causing any steric clashes. In the “down” conformation, the RBDs were tightly packed against the top of the S2 subunit, preventing binding by ACE2 (Gui et al., 2017). Subsequent experiments have corroborated this phenomenon and similar dynamics have been observed in MERS-CoV S, SARS-CoV-2 S and in alphacoronavirus S proteins (Jones et al., 2019; Kirchdoerfer et al., 2018; Pallesen et al., 2017; Walls, 2020; Wrapp et al., 2020; Yuan et al., 2017). Due to the relatively low abundance of particles that can be observed by cryo-EM with three RBDs in the up conformation, it is thought that this conformation may correspond to an energetically unstable state (Kirchdoerfer et al., 2018; Pallesen et al., 2017). These observations led to the hypothesis that the CoV RBDs may be acting as molecular ratchets, wherein a receptor-binding event would trap the RBD in the less stable up conformation, leading to gradual destabilization until S is finally triggered to initiate membrane fusion. Recent experiments characterizing RBD-directed anti-SARS-CoV-1 antibodies that trap the SARS-CoV-1 RBD in the up conformation and lead to destabilization of the prefusion spike have lent support to this hypothesis (Walls et al., 2019).

Numerous anti-SARS-CoV-1 RBD and anti-MERS-CoV RBD antibodies have been reported and their mechanisms of neutralization can be attributed to the occlusion of the receptor-binding site and to trapping the RBD in the unstable up conformation, effectively acting as a receptor mimic that triggers a premature transition from the prefusion-to-postfusion conformation (Hwang et al., 2006; Walls et al., 2019; Wang et al., 2018; Wang et al., 2015). Heavy chain-only antibodies (HCAbs), present in camelids, contain a single variable domain (VHH) instead of two variable domains (VH and VL) that make up the equivalent antigen-binding fragment (Fab) of conventional IgG antibodies (Hamers-Casterman et al., 1993). This single variable domain, in the absence of an effector domain, is referred to as a single-domain antibody, VHH or Nanobody® and typically can acquire affinities and specificities for antigens comparable to conventional antibodies. VHHs can easily be constructed into multivalent formats and are known to have enhanced thermal stability and chemostability compared to most antibodies (De Vlieger et al., 2018; Dumoulin et al., 2002; Govaert et al., 2012; Laursen et al., 2018; van der Linden et al., 1999). Their advantageous biophysical properties have led to the evaluation of several VHHs as therapeutics against common respiratory pathogens, such as respiratory syncytial virus (RSV) (Detalle et al., 2016; Rossey et al., 2017). The use of VHHs as biologics in the context of a respiratory infection is a particularly attractive application, since the highly stable VHHs can be nebulized and administered via an inhaler directly to the site of infection (Respaud et al., 2015). Moreover, due to their stability after prolonged storage, VHHs could be stockpiled as therapeutic treatment options in case of an epidemic. Although therapeutics against MERS-CoV and SARS-CoV-2 are sorely needed, the feasibility of using VHHs for this purpose has not yet been adequately explored. Several MERS-CoV S-directed VHHs have been reported, but their epitopes remain largely undefined, other than being classified as RBD-directed (Stalin Raj et al., 2018; Zhao et al., 2018).

Here we report the isolation of two potently neutralizing VHHs directed against the SARS-CoV-1 and MERS-CoV RBDs. These VHHs were elicited in response to immunization of a llama with prefusion-stabilized SARS-CoV-1 and MERS-CoV S proteins. We solved the crystal structures of these two VHHs in complex with their respective viral epitopes and determined that their mechanisms of neutralization were occlusion of the receptor binding interface and trapping of the RBDs in the up conformation. We also show that the SARS-CoV-1 RBD-directed VHH exhibits cross-reactivity against the SARS-CoV-2 RBD and is capable of blocking the receptor-binding interface. After engineering this VHH into a bivalent Fc-fusion, we show that this cross-reactive VHH is also capable of potently neutralizing SARS-CoV-2 S pseudoviruses. In addition, we demonstrate that the VHH-Fc fusion can be produced at high yields in an industry-standard CHO cell system, suggesting that it merits further investigation as a potential therapeutic for the ongoing COVID-19 pandemic.

## RESULTS

### Isolation of betacoronavirus S-directed VHHs

Our initial aim was to isolate VHHs that could potently neutralize MERS-CoV and SARS-CoV-1. Therefore, a llama was sequentially immunized subcutaneously twice with SARS-CoV-1 S protein, twice with MERS-CoV S protein, once again with SARS-CoV-1 S and finally with both SARS-CoV-1 and MERS-CoV S protein (**S. Figure 1A**). To obtain VHHs directed against these spike proteins, two consecutive rounds of panning were performed using either SARS-CoV-1 S or MERS-CoV S protein. Positive clones were sequenced and multiple sequence alignment and phylogenetic analysis using the neighbor-joining method revealed that seven unique MERS-CoV S and five unique SARS-CoV-1 S VHHs were isolated (**S. Figure 1B**). These VHHs and an irrelevant control (RSV F-VHH, directed against the F protein of human respiratory syncytial virus) were subsequently expressed in *Pichia pastoris* and purified from the yeast medium (Rossey et al., 2017). The binding of the purified VHHs to prefusion-stabilized MERS-CoV S and SARS-CoV-1 S was confirmed by ELISA (**S. Figure 1C**). Four clones (MERS VHH-55, −12, −34 and −40), obtained after panning on MERS-CoV S protein, bound with high affinity to prefusion stabilized MERS-CoV S, whereas the affinity of VHH-2, −20 and −15 was 100- to 1000-fold lower. Of the five clones isolated after panning on SARS-CoV-1 S protein, three VHH clones (SARS VHH-72, −1 and −6) interacted strongly with prefusion stabilized SARS-CoV-1 S protein.

**Figure 1:**
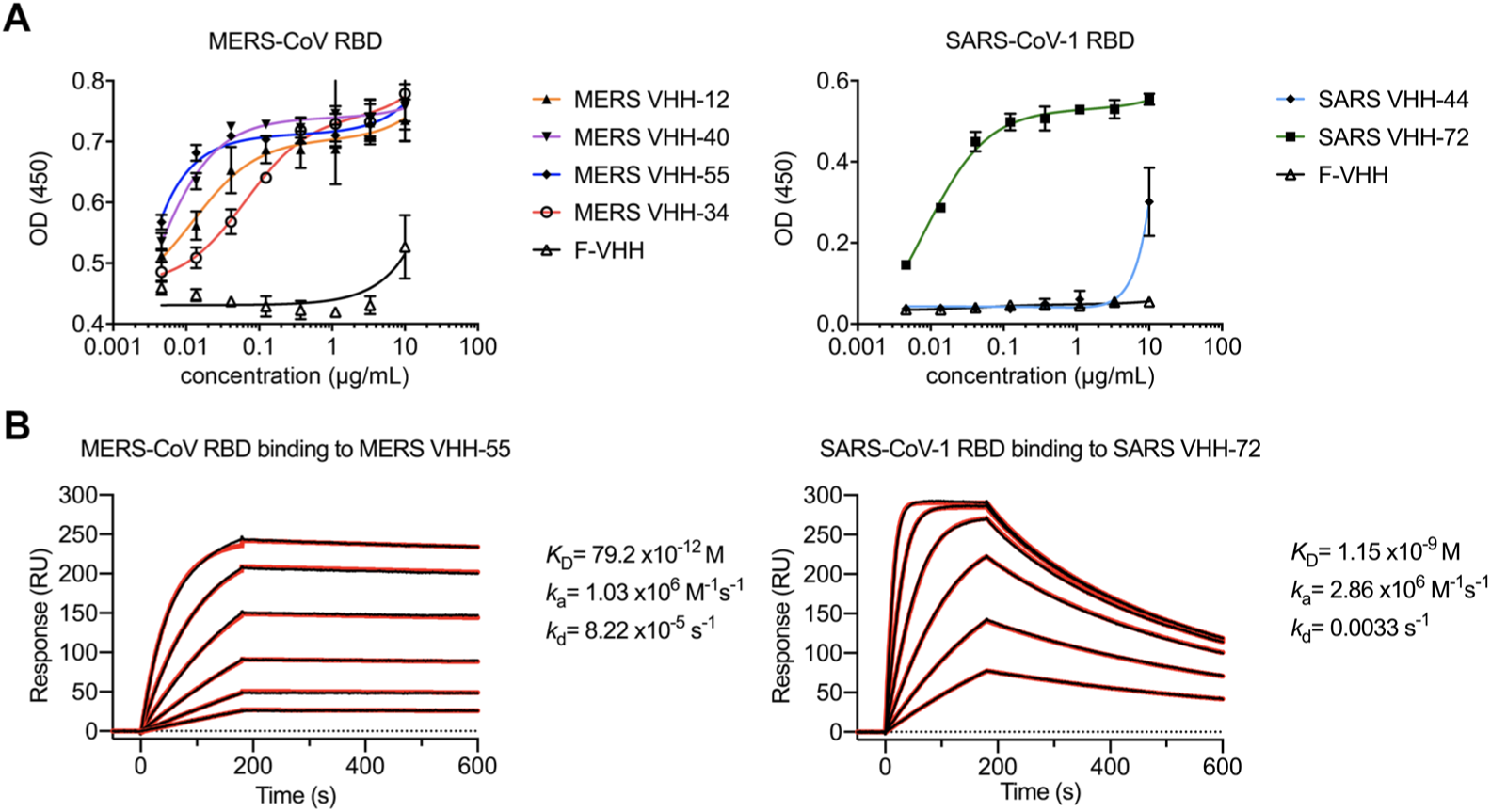
Epitope determination and biophysical characterization of MERS VHH-55 and SARS VHH-72. **A**) Reactivity of MERS-CoV and SARS-CoV RBD-directed VHHs against the MERS-CoV and SARS-CoV-1 RBD, respectively. A VHH against an irrelevant antigen (F-VHH) was included as a control. **B**) SPR sensorgrams showing binding between the MERS-CoV RBD and MERS VHH-55 (*left*) and SARS-CoV-1 RBD and SARS VHH-72 (*right*). Binding curves are colored black and fit of the data to a 1:1 binding model is colored red.

### VHHs neutralize coronavirus S pseudotyped viruses

To assess the antiviral activity of the MERS-CoV and SARS-CoV S-directed VHHs, *in vitro* neutralization assays using MERS-CoV England1 S and SARS-CoV-1 Urbani S pseudotyped lentiviruses were performed. The high affinity MERS VHH-55, −12, −34 and −40 neutralized MERS-CoV S pseudotyped virus with IC_50_ values ranging from 0.014 to 2.9 µg/mL (0.9 nM to 193.3 nM), while no inhibition was observed for the lower affinity MERS-CoV or SARS-CoV-1 specific VHHs (**Table 1**). SARS VHH-72 and −44 were able to neutralize lentiviruses pseudotyped with SARS-CoV-1 S with an IC_50_ value of 0.14 (9 nM) and 5.5 µg/mL (355 nM), respectively. No binding was observed for SARS VHH-44 to prefusion stabilized SARS-CoV-1 S protein in the ELISA assay. Sequence analysis revealed that the neutralizing MERS-CoV specific VHHs −12, −40 and −55 have highly similar complementarity-determining regions (CDRs), indicating that they likely bind to the same epitope (**S. Figure 2**). In contrast, the CDRs from the SARS-CoV S-specific VHHs −44 and −72 are highly divergent.

**Figure 2:**
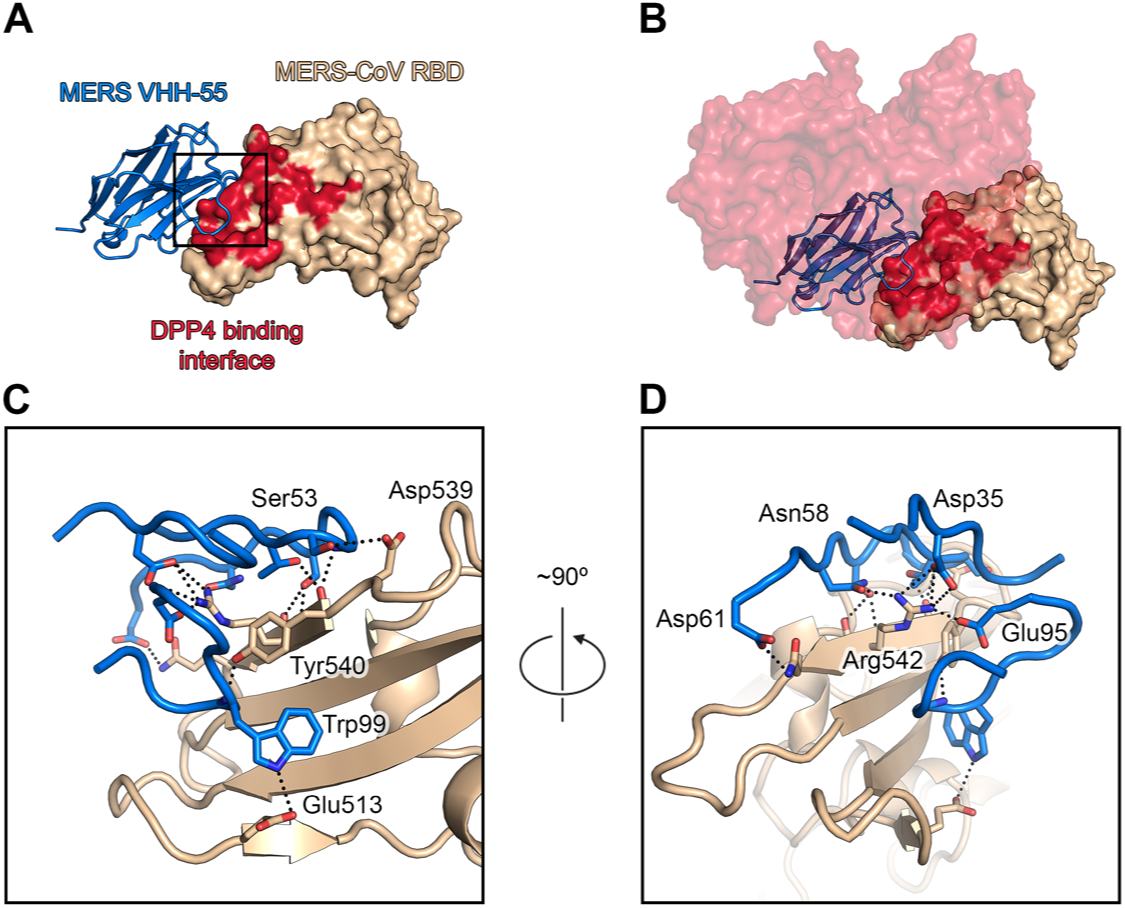
The crystal structure of MERS VHH-55 bound to the MERS-CoV RBD. **A**) MERS VHH-55 is shown as blue ribbons and the MERS-CoV RBD is shown as a tan-colored molecular surface. The DPP4 binding interface on the MERS-CoV RBD is colored red. **B**) The structure of DPP4 bound to the MERS-CoV RBD (PDB: 4L72) is aligned to the crystal structure of MERS VHH-55 bound to the MERS-CoV RBD. A single monomer of DPP4 is shown as a red, transparent molecular surface. **C**) A zoomed-in view of the panel from **2A**, with the MERS-CoV RBD now displayed as tan-colored ribbons. Residues that form interactions are shown as sticks, with nitrogen atoms colored dark blue and oxygen atoms colored red. Hydrogen-bonds and salt bridges between MERS VHH-55 and the MERS-CoV RBD are shown as black dots. **D**) The same view from **2C** has been turned by approximately 90° to show additional contacts. Residues that form interactions are shown as sticks, with nitrogen atoms colored dark blue and oxygen atoms colored red. Hydrogen-bonds and salt bridges between MERS VHH-55 and the MERS-CoV RBD are shown as black dots.

**Table 1:** MERS-CoV and SARS-CoV-1 pseudovirus neutralization data.

|  | Neutralization IC<sub>50</sub>(μg/mL) |  |
| --- | --- | --- |
|  | MERS-CoV<br>England 1 | SARS-CoV-1<br>Urbani |
| D12 mAb (ctrl) | 0.01996 | >10 |
| MERS VHH-2 | >10 | >10 |
| MERS VHH-12 | 0.13 | >10 |
| MERS VHH-15 | >10 | >10 |
| MERS VHH-20 | >10 | >10 |
| MERS VHH-34 | 2.9 | >10 |
| MERS VHH-40 | 0.034 | >10 |
| MERS VHH-55 | 0.014 | >10 |
| SARS VHH-1 | >10 | >10 |
| SARS VHH-6 | >10 | >10 |
| SARS VHH-35 | >10 | >10 |
| SARS VHH-44 | >10 | 5.5 |
| SARS VHH-72 | >10 | 0.14 |

### Epitope mapping of betacoronavirus S-directed VHHs

To map the epitopes targeted by the neutralizing VHHs, we tested binding to recombinant MERS-CoV S1, RBD, NTD and SARS-CoV-1 RBD and NTD by ELISA (**Figure 1A and S**. **Figure 3**).The MERS-CoV S-specific VHHs strongly bound to MERS-CoV S1 and RBD in a concentration-dependent manner, and no binding to the MERS-CoV NTD was observed. Similarly, strong binding of SARS VHH-72 to the SARS-CoV-1 RBD protein but not the SARS-CoV-1 NTD protein was observed. Again, no binding of SARS VHH-44 to either the SARS-CoV-1 RBD or NTD protein was detected. These data demonstrate that the neutralizing VHHs SARS VHH-72 and MERS VHH-55 target the RBDs. Based on the specificity and potent neutralizing capacity of SARS VHH-72 and MERS VHH-55, we measured the affinities of these VHHs by immobilizing recombinantly expressed VHH to an SPR sensorchip and determined the binding kinetics for their respective RBDs. We found that both of these VHHs bound to their targets with high affinity. SARS VHH-72 bound to its target with an affinity of 1.2 nM and MERS VHH-55 bound to its target with an affinity of 79.2 pM, in part due to a very slow off-rate constant (*k*_d_ = 8.2 x10^-5^ s^-1^) (**Figure 1B**).

**Figure 3:**
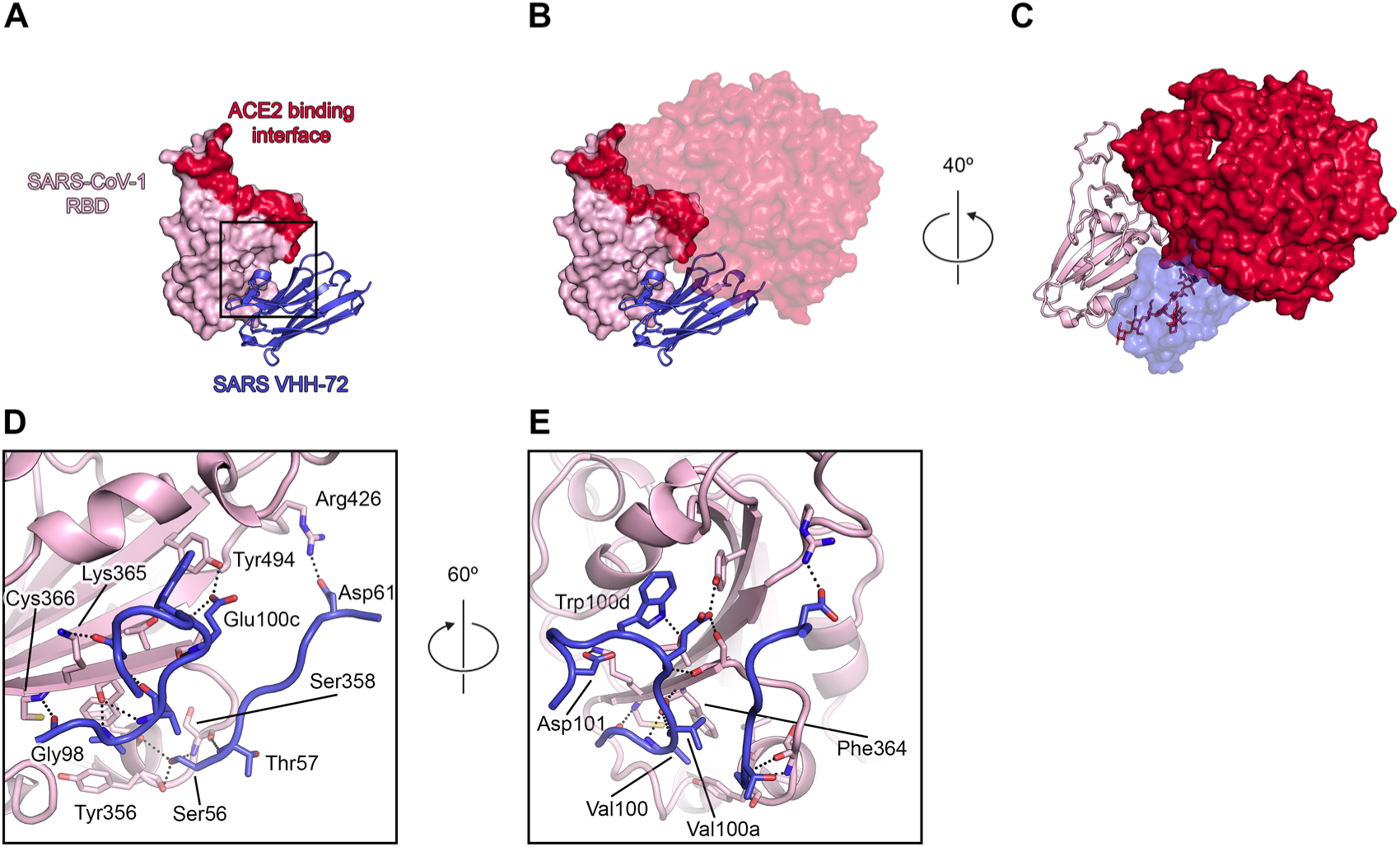
The crystal structure of SARS VHH-72 bound to the SARS-CoV-1 RBD. **A**) SARS VHH-72 is shown as dark blue ribbons and the SARS-CoV-1 RBD is shown as a pink-colored molecular surface. The ACE2 binding interface on the SARS-CoV-1 RBD is colored red. **B**) The structure of ACE2 bound to the SARS-CoV-1 RBD (PDB: 2AJF) is aligned to the crystal structure of SARS VHH-72 bound to the SARS-CoV-1 RBD. ACE2 is shown as a red, transparent molecular surface. **C**) A simulated *N*-linked glycan containing an energy-minimized trimannosyl core (derived from PDB ID: 1HD4) is modeled as red sticks, coming from Asn322 in ACE2. ACE2 is shown as a red molecular surface, the SARS-CoV-1 RBD is shown as pink ribbons and SARS VHH-72 is shown as a dark blue, transparent molecular surface. **D**) A zoomed-in view of the panel from **3A** is shown, with the SARS-CoV-1 RBD now displayed as pink-colored ribbons. Residues that form interactions are shown as sticks, with nitrogen atoms colored dark blue and oxygen atoms colored red. Hydrogen bonds and salt bridges between SARS VHH-72 and the SARS-CoV-1 RBD are shown as black dots. **E**) The same view from **3D** has been turned by 60° to show additional contacts. Residues that form interactions are shown as sticks, with nitrogen atoms colored dark blue and oxygen atoms colored red. Interactions between SARS VHH-72 and the SARS-CoV-1 RBD are shown as black dots.

### Structural basis of VHH interaction with RBDs

To investigate the molecular determinants that mediate potent neutralization and high-affinity binding by MERS VHH-55, we solved the crystal structure of MERS VHH-55 bound to the MERS-CoV RBD. Crystals grew in space group *C*222_1_ and diffracted X-rays to a resolution of 3.4 Å. After determining a molecular replacement solution and iterative building and refinement, our structure reached an R_work_/R_free_ of 21.4%/26.8% (**Table 2**). The asymmetric unit of this crystal contained eight copies of the MERS VHH-55 + MERS-CoV RBD complex and had a solvent content of ∼58%. The electron density allowed unambiguous definition of the interface between the RBD and VHH, with the three CDRs forming extensive binding contacts with the RBD, burying 716 Å^2^ of surface area by pinching the RBD between the CDR2 and CDR3. The CDR3 of MERS VHH-55 is looped over the DPP4-binding interface, occluding DPP4 from productively engaging the MERS-CoV RBD (**Figure 2A-B**).

**Table 2:**
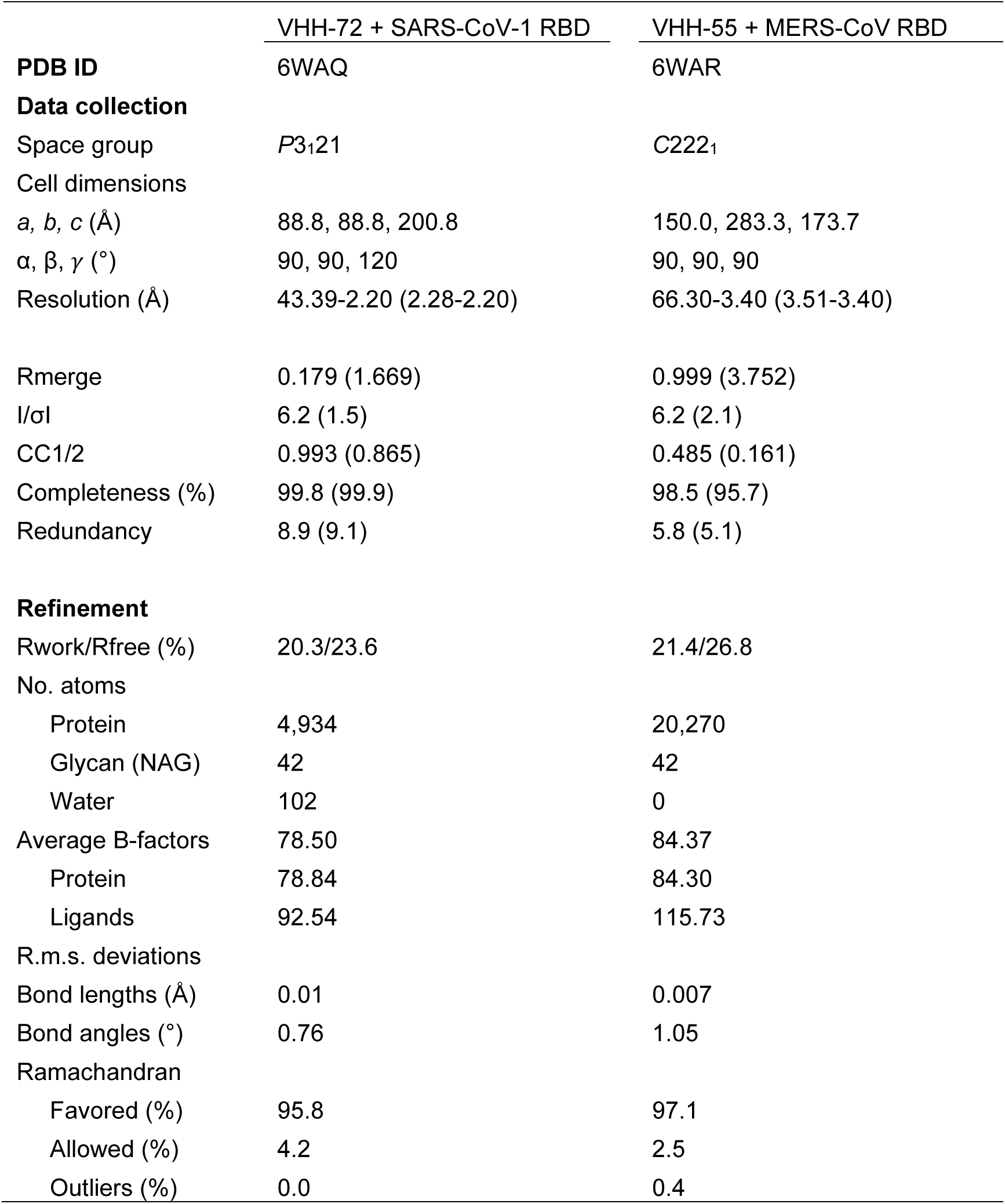
X-ray data collection and refinement statistics

There are numerous contacts between the CDRs of MERS VHH-55 and the MERS-CoV RBD, and the majority of these are confined to CDRs 2 and 3 (**Figure 2C-D**). The sole interaction from the MERS VHH-55 CDR1 comes from Asp35 forming a salt bridge with Arg542 from the RBD. CDR2 forms hydrogen bonds using Ser53 and Asp61 to engage RBD residues Asp539 and Gln544. Furthermore, Asn58 from the CDR2 also engages Arg542 from the RBD. Trp99 from the MERS VHH-55 CDR3 forms a salt bridge with Glu513 via the nitrogen from its pyrrole ring. Finally, Glu95 from the MERS VHH-55 CDR3 also forms a salt bridge with Arg542 from the MERS-CoV RBD, suggesting that Arg542 plays a critical role in MERS VHH-55 binding since it is productively engaged by residues from all three CDRs. This arginine has also been implicated in binding to the MERS-CoV receptor DPP4, and has previously been identified as one of the twelve highly conserved amino acids that is crucial for high-affinity receptor engagement (**S. Figure 4A**) (Wang et al., 2013; Wang et al., 2014).

**Figure 4:**
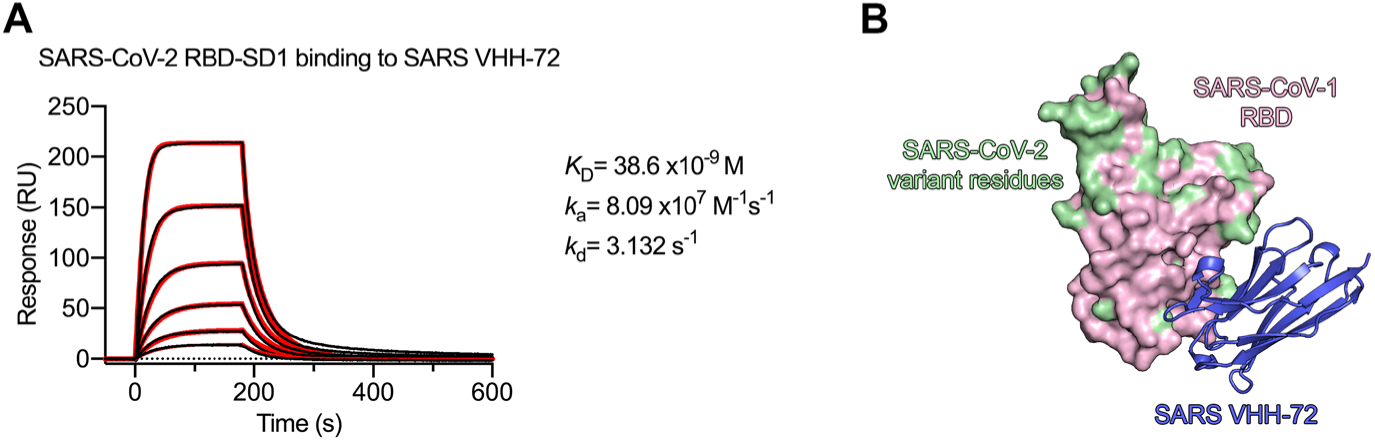
SARS VHH-72 is cross-reactive against SARS-CoV-2. **A**) An SPR sensorgram measuring the binding of SARS VHH-72 to the SARS-CoV-2 RBD-SD1. Binding curves are colored black and fit of the data to a 1:1 binding model is colored red. **B**) The crystal structure of SARS VHH-72 bound to the SARS-CoV-1 RBD is shown with SARS VHH-72 as dark blue ribbons and the RBD as a pink molecular surface. Amino acids that vary between SARS-CoV-1 and SARS-CoV-2 are colored green.

In addition to forming a salt bridge with Glu513 from the MERS-CoV RBD, Trp99 of the MERS VHH-55 CDR3 is positioned near a hydrophobic patch formed by Phe506 (**S. Figure 4B**). This amino acid exhibits natural sequence variation in several MERS-CoV strains, such that a Leu is occasionally observed at this position. To evaluate the extent to which this substitution may impact MERS VHH-55 binding, we generated a F506L substitution and measured binding by SPR (**S. Figure 4C**). Surprisingly, this substitution resulted in a ∼200-fold reduction in MERS VHH-55 binding affinity. Despite this substantial reduction, the affinity of MERS VHH-55 to MERS-CoV RBD F506L remains high, with a *K*_D_ = 16.5 nM. Other than the variability that is observed at position 506 of the MERS-CoV RBD, the rest of the MERS VHH-55 epitope is highly conserved across the 863 strains that are curated in the MERS-CoV Virus Variation database (**S. Figure 4A**). This high degree of epitope conservation suggests that VHH-55 would broadly recognize MERS-CoV strains.

We also sought to discover the molecular determinants of binding between SARS VHH-72 and the SARS-CoV-1 RBD by determining the crystal structure of this complex. Crystals grew in space group *P*3_1_21 and diffracted X-rays to a resolution of 2.2 Å. We obtained a molecular replacement solution and refined the structure to an R_work_/R_free_ of 20.3%/23.6% through iterative building and refinement (**Table 2**). Our structure reveals that CDRs 2 and 3 contribute to the majority of the 834 Å^2^ of buried surface area at the binding interface (**Figure 3A**). This interface does not, however, overlap with the ACE2 binding interface. Rather, ACE2 would clash with the CDR-distal framework of SARS VHH-72 (**Figure 3B**). This clash would only be enhanced by the presence of an *N*-linked glycan at Asn322 of ACE2, which is already located within the VHH framework when the receptor-bound RBD is aligned to the VHH-bound RBD (**Figure 3C**).

SARS VHH-72 binds to the SARS-CoV-1 RBD by forming an extensive hydrogen-bond network via its CDRs 2 and 3 (**Figure3D-E**). Ser56 from the CDR2 simultaneously forms hydrogen bonds with the peptide backbone of three residues from the SARS-CoV-1 RBD: Leu355, Tyr356 and Ser358. The peptide backbone of Ser358 also forms a hydrogen bond with the backbone of neighboring Thr57 from the CDR2. A salt bridge formed between CDR2 residue Asp61 and RBD residue Arg426 tethers the C-terminal end of the CDR2 to the RBD. The N-terminus of the CDR3 forms a short β-strand that pairs with a β-strand from the SARS-CoV-1 RBD to bridge the interface between these two molecules. This interaction is mediated by backbone hydrogen bonds from CDR3 residues Gly98, Val100 and Val100a to RBD residues Cys366 and Phe364. Glu100c from the CDR3 forms hydrogen bonds with the sidechain hydroxyls from both Ser362 and Tyr494 from the SARS-CoV-1 RBD. The neighboring CDR3 residue also engages in a sidechain-specific interaction by forming a salt bridge between the pyrrole nitrogen of Trp100d and the hydroxyl group from RBD residue Thr363. Asp101 is involved in the most C-terminal interaction from the CDR3 by forming a salt bridge with RBD residue Lys365. The extensive interactions formed between CDRs 2 and 3 of SARS VHH-72 and the SARS-CoV-1 RBD explain the high-affinity binding that we observed between these molecules.

### SARS VHH-72 is cross-reactive against WIV1-CoV and SARS-CoV-2

Analysis of 10 available SARS-CoV-1 strain sequences revealed a high degree of conservation in the residues that make up the SARS VHH-72 epitope, prompting us to explore the breadth of SARS VHH-72 binding (**S. Figure 5A**). WIV1-CoV is a betacoronavirus found in bats that is closely related to SARS-CoV-1 and also utilizes ACE2 as a host-cell receptor (Ge et al., 2013). Due to the relatively high degree of sequence conservation between SARS-CoV and WIV1-CoV, we expressed the WIV1-CoV RBD and measured binding to SARS VHH-72 by SPR (**S. Figure 5B**). SARS VHH-72 also exhibits high-affinity binding to the WIV1-CoV RBD (7.4 nM), demonstrating that it is cross-reactive between these two closely related coronaviruses (**S. Figure 5C**).

**Figure 5:**
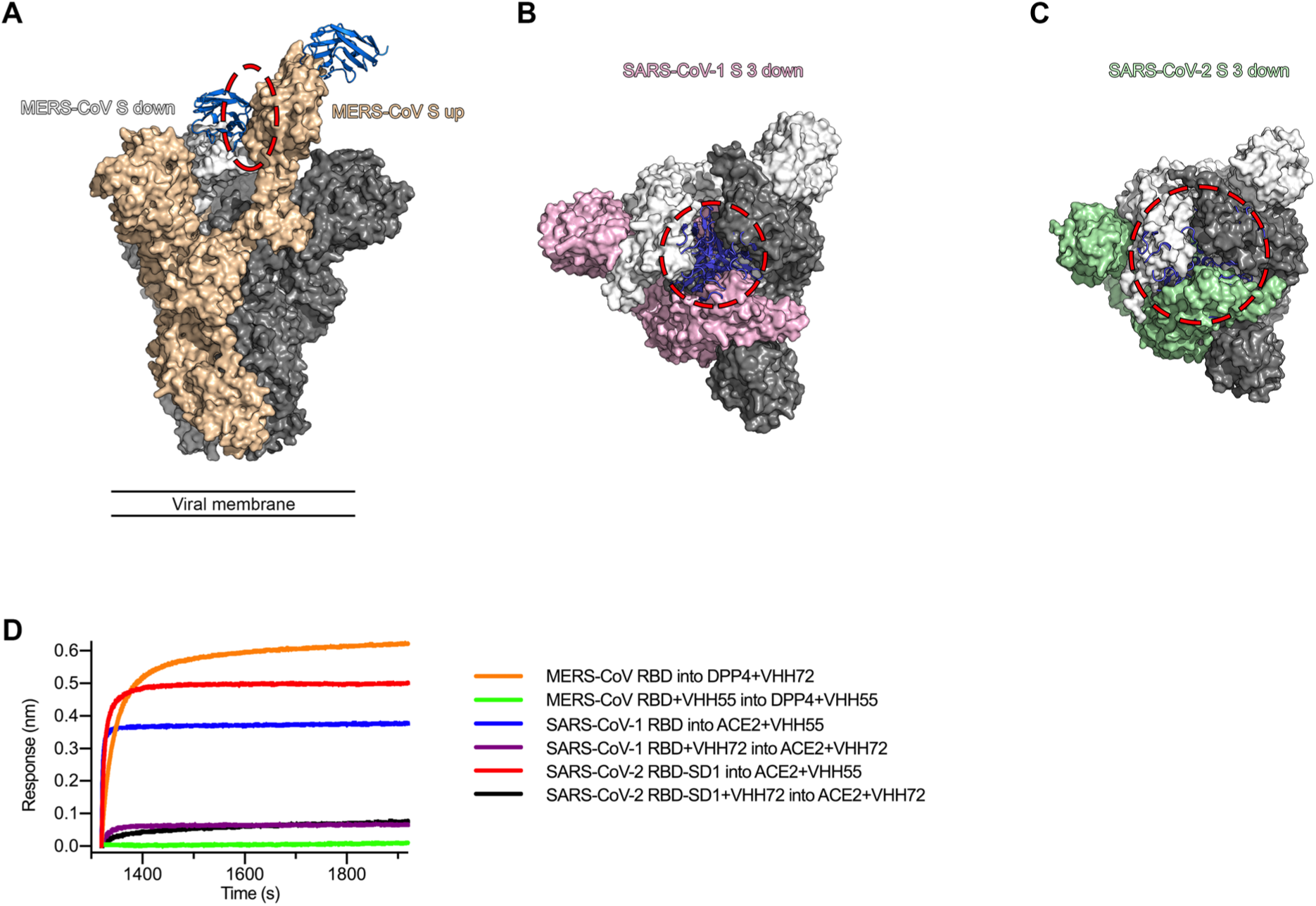
Neutralizing mechanisms of MERS VHH-55 and SARS VHH-72. **A**) The MERS-CoV spike (PDB ID: 5W9H) is shown as a molecular surface, with each monomer colored either white, gray or tan. The tan and white monomers are bound by MERS VHH-55, shown as blue ribbons. The clash between MERS VHH-55 bound to the white monomer and the neighboring tan RBD is highlighted by the red ellipse. **B**) The SARS-CoV-1 spike (PDB ID: 5X58) is shown as a molecular surface, with each protomer colored either white, gray or pink. Every monomer is bound by a copy of SARS VHH-72, shown as dark blue ribbons. The clashes between copies of SARS VHH-72 and the two neighboring spike monomers are highlighted by the red circle. **C**) The SARS-CoV-2 spike (PDB ID: 6VXX) is shown as a molecular surface, with each protomer colored either white, gray or green. Every monomer is bound by a copy of SARS VHH-72, shown as dark blue ribbons. The clashes between copies of SARS VHH-72 and the two neighboring spike monomers are highlighted by the red circle. The SARS-CoV-2 trimer appears smaller than SARS-CoV-1 S due to the absence of flexible NTD-distal loops which could not be built during cryo-EM analysis. **D)** CoV VHHs prevent MERS-CoV RBD, SARS-CoV-1 RBD and SARS-CoV-2 RBD-SD1 from interacting with their receptors. The results of the BLI-based receptor-blocking experiment are shown. The legend lists the immobilized RBDs and the VHHs or receptors that correspond to each curve.

Based on the high degree of structural homology that has been reported between SARS-CoV-1 S and SARS-CoV-2 S (Walls, 2020; Wrapp et al., 2020), we also tested SARS VHH-72 for cross-reactivity against the SARS-CoV-2 RBD-SD1 by SPR (**Figure 4**). The binding affinity of SARS VHH-72 for the SARS-CoV-2 RBD-SD1 was ∼39 nM. This diminished binding affinity, compared to the binding of SARS VHH-72 to SARS-CoV-1 RBD, can primarily be attributed to an increase in the dissociation rate constant of this interaction (**Figure 4A**). The only variant residue on the SARS-CoV-1 RBD that makes direct contact with SARS VHH-72 is Arg426, which is Asn439 in the SARS-CoV-2 RBD (**Figure 3C**).

### VHHs disrupt RBD dynamics and receptor-binding

As stated previously, the RBDs of MERS-CoV S, SARS-CoV-1 S and SARS-CoV-2 S undergo dynamic conformational rearrangements that alternately mask and present their receptor-binding interfaces and potential neutralizing epitopes to host molecules. By aligning the crystal structures of MERS VHH-55 and SARS VHH-72 bound to their respective targets to the full-length, cryo-EM structures of the MERS-CoV, SARS-CoV-1 and SARS-CoV-2 spike proteins, we can begin to understand how these molecules might function in the context of these dynamic rearrangements. When the MERS-CoV RBDs are all in the down conformation or all in the up conformation, MERS VHH-55 would be able to bind all three of the protomers making up the functional spike trimer without forming any clashes. However, if a down protomer was bound by MERS VHH-55 and the neighboring protomer sampled the up conformation, this RBD would then be trapped in this state by the presence of the neighboring MERS VHH-55 molecule (**Figure 5A**). This conformational trapping would be even more pronounced upon SARS VHH-72 binding to the full-length SARS-CoV-1 S protein or the full-length SARS-CoV-2 S protein.

Due to the binding angle of SARS VHH-72, when a bound SARS-CoV-1 or SARS-CoV-2 RBD samples the down conformation, it would form dramatic clashes with the S2 fusion subunit, regardless of the conformations of the neighboring RBDs (**Figure 5B-C**). Therefore, once a single SARS VHH-72 binding event took place, the bound protomer would be trapped in the up conformation until either SARS VHH-72 was released or until the S protein was triggered to undergo the prefusion-to-postfusion transition. Based on the binding angles of MERS VHH-55 and SARS VHH-72, we can conclude that these molecules would likely disrupt the RBD dynamics in the context of a full-length S protein by trapping the up conformation. Because this up conformation is unstable and leads to S protein triggering, it is possible that this conformational trapping may at least partially contribute to the neutralization mechanisms of these VHHs.

Based on our structural analysis, we hypothesized that another mechanism by which both MERS VHH-55 and SARS VHH-72 neutralize their respective viral targets is by blocking the interaction between the RBDs and their host-cell receptors. To test this hypothesis, we performed a BLI-based assay in which the SARS-CoV-1, SARS-CoV-2 and MERS-CoV RBDs were immobilized to biosensor tips, dipped into VHHs and then dipped into wells containing the recombinant, soluble host cell receptors. We found that when tips coated in the MERS-CoV RBD were dipped into MERS VHH55 before being dipped into DPP4, there was no increase in response that could be attributed to receptor binding. When tips coated with the MERS-CoV RBD were dipped into SARS VHH-72 and then DPP4, a robust response signal was observed, as expected. Similar results were observed when the analogous experiments were performed using the SARS-CoV-1 or SARS-CoV-2 RBDs, SARS VHH-72 and ACE2 (**Figure 5D**). These results support our structural analysis that both MERS VHH-55 and SARS VHH-72 are capable of neutralizing their respective viral targets by directly preventing host-cell receptor binding.

### Bivalent SARS VHH-72 is capable of potently neutralizing SARS-CoV-2 S pseudoviruses

Despite the relatively high-affinity binding that was observed by SPR between SARS VHH-72 and the SARS-CoV-2 RBD, this binding could not be detected by ELISA nor was SARS VHH-72 capable of neutralizing SARS-CoV-2 S VSV pseudoviruses, possibly due to the high off-rate constant, whereas SARS-CoV-1 pseudotypes were readily neutralized (**Figure 6A-D**). In an attempt to overcome this rapid dissociation, we engineered two bivalent variants of SARS VHH-72. These included a tail-to-head fusion of two SARS VHH-72 molecules connected by a (GGGGS)_3_ linker (VHH-72-VHH-72) and a genetic fusion of SARS VHH-72 to the Fc domain of human IgG1 (VHH-72-Fc) (**S. Figure 6A-C**). These bivalent SARS VHH-72 constructs were able to bind to both prefusion SARS-CoV-1 S and SARS-CoV-2 RBD-SD1 as demonstrated by ELISA and by a dose-dependent reduction in the binding of SARS-CoV-2 RBD-SD1 to Vero E6 cells (**Figure 6C-D and S. Figure 6B-C**). We could also detect binding of both of these constructs to full length SARS-CoV-1 S and SARS-CoV-2 S expressed on the surface of mammalian cells (**S. Figure 6D-E**). Supernatants of HEK 293S cells transiently transfected with VHH-72-Fc exhibited neutralizing activity against both SARS-CoV-1 and SARS-CoV-2 S VSV pseudoviruses in the same assay which showed no such cross-reactive neutralization for monovalent SARS VHH-72 (**Figure 6E-F**). A BLI experiment measuring binding of VHH-72-Fc to immobilized SARS-CoV-2 RBD-SD1 further confirmed that bivalency was able to compensate for the high off-rate constant of the monomer (**Figure 7A**). Furthermore, cross-neutralizing VHH-72-Fc construct reached expression levels of ∼300 mg/L in ExpiCHO cells (**Figure 7B**). Using VHH-72-Fc purified from ExpiCHO cells and a SARS-CoV-2 S pseudotyped VSV with a luciferase reporter, we evaluated the neutralization capacity of VHH-72-Fc and found that it was able to neutralize pseudovirus with an IC_50_ of approximately 0.2 μg/mL (**Figure 7C**).

**Figure 6:**
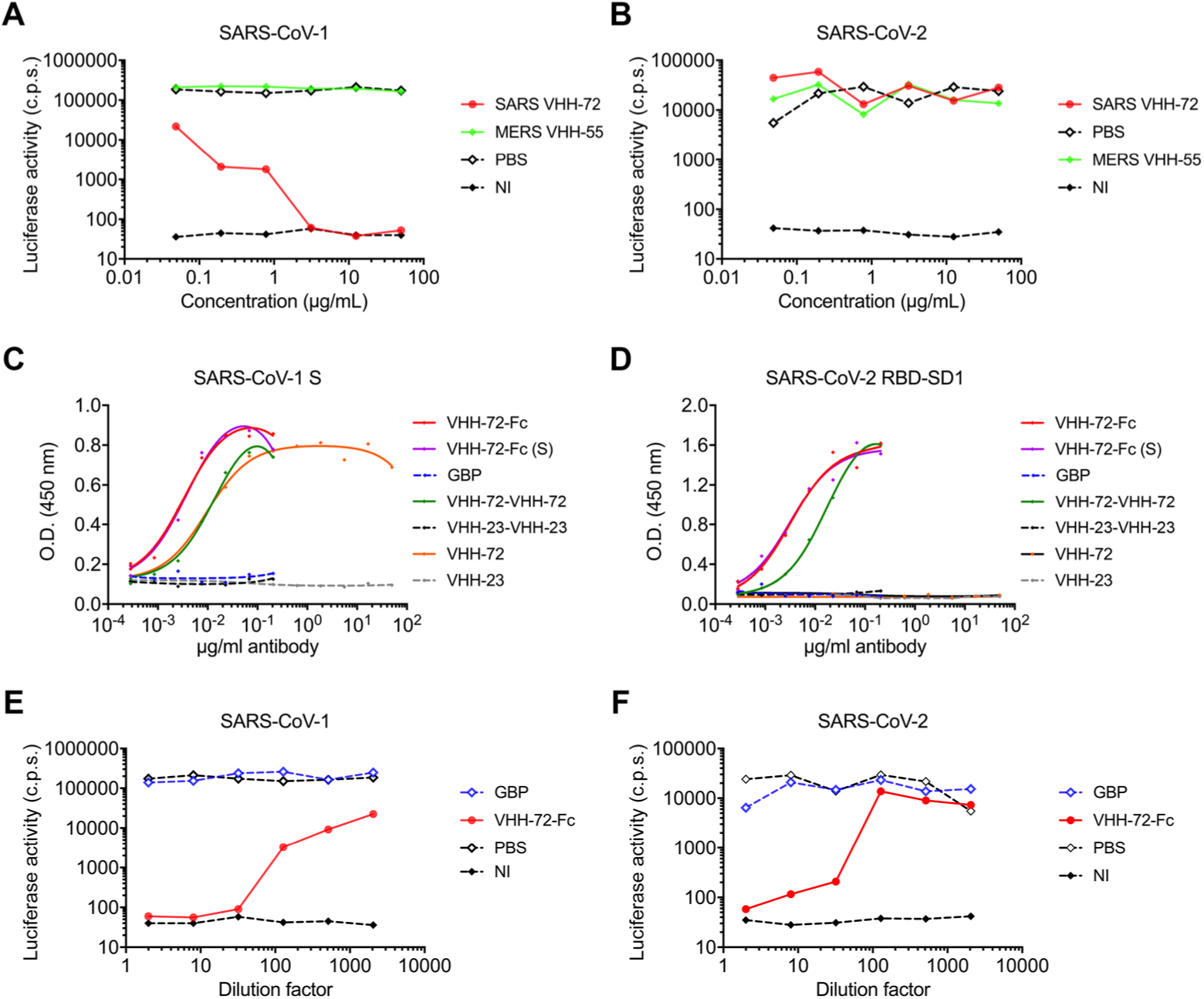
Bivalency overcomes the high off-rate constant of SARS VHH-72. **A**) SARS-CoV-1 S and **B**) SARS-CoV-2 S VSV pseudoviruses were used to evaluate the neutralization capacity of SARS VHH-72. MERS VHH-55 and PBS were included as negative controls. Luciferase activity is reported in counts per second (c.p.s.). NI cells were not infected. **C**) Binding of bivalent VHHs was tested by ELISA against SARS-CoV-1 S and **D**) SARS-CoV-2 RBD-SD1. VHH-72-Fc refers to SARS VHH-72 fused to a human IgG1 Fc domain by a GS(GGGGS)_2_ linker. VHH-72-Fc (S) is the same Fc fusion with a GS, rather than a GS(GGGGS)_2_, linker. GBP is an irrelevant GFP-binding protein. VHH-72-VHH-72 refers to the tail-to-head construct with two SARS VHH-72 proteins connected by a (GGGGS)_3_ linker. VHH-23-VHH-23 refers to the two irrelevant VHHs linked via the same (GGGGS)_3_ linker. **E**) SARS-CoV-1 S and **F**) SARS-CoV-2 S pseudoviruses were used to evaluate the neutralization capacity of bivalent VHH-72-Fc. GBP and PBS were included as negative controls. NI cells were not infected.

**Figure 7:**
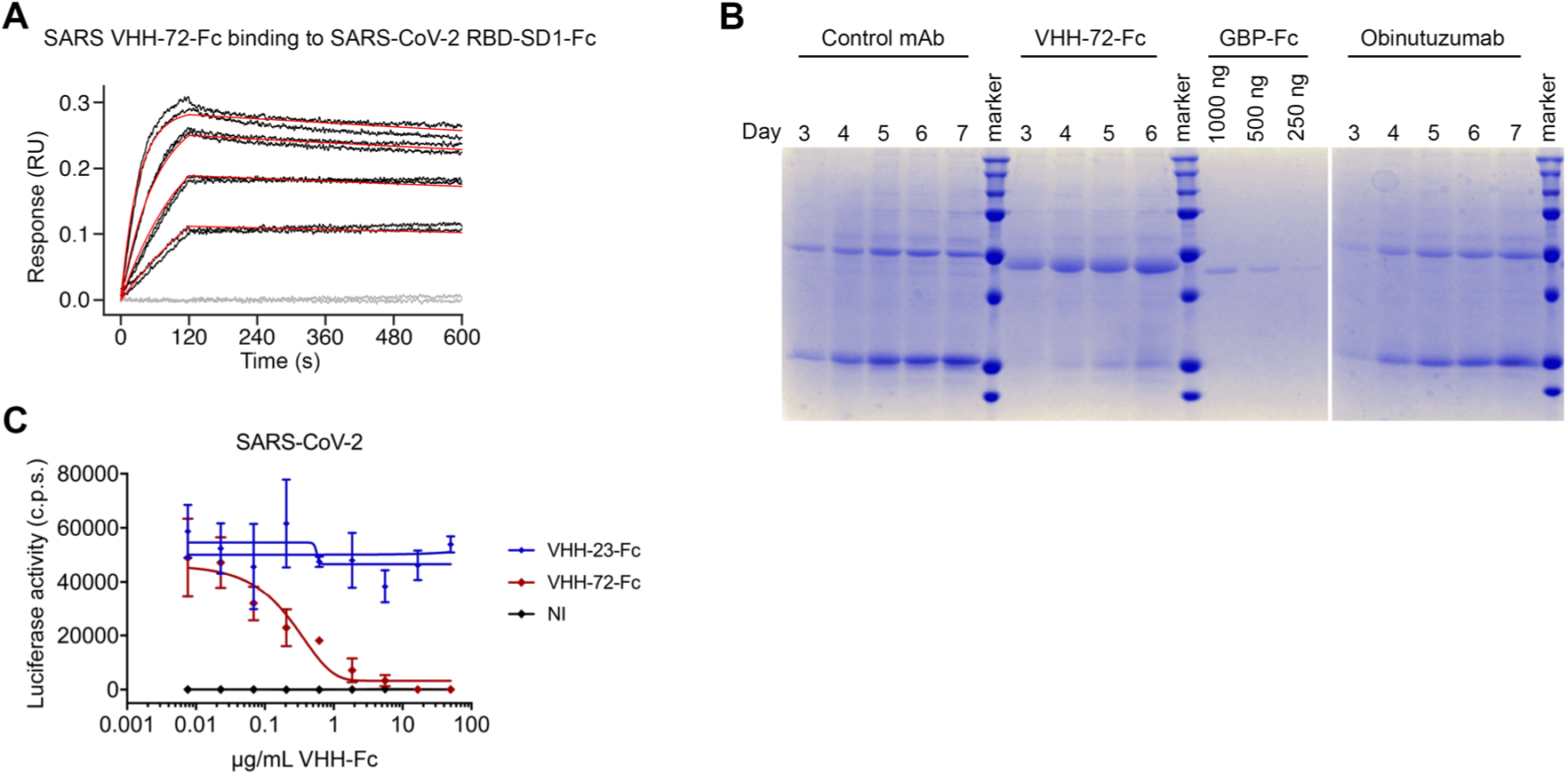
VHH-72-Fc neutralizes SARS-CoV-2 S pseudoviruses. **A**) BLI sensorgram measuring apparent binding affinity of VHH-72-Fc to immobilized SARS-CoV-2 RBD-SD1. Binding curves are colored black, buffer-only blanks are colored gray and the fit of the data to a 1:1 binding curve is colored red. **B**) Time course analysis of VHH-72-Fc expression in ExpiCHO cells. Cell culture supernatants of transiently transfected ExpiCHO cells were removed on days 3-7 after transfection (or until cell viability dropped below 75%), as indicated. Two control mAbs were included for comparison, along with the indicated amounts of purified GBP-Fc as a loading control. **C**) SARS-CoV-2 S pseudotyped VSV neutralization assay. Monolayers of Vero E6 cells were infected with pseudoviruses that had been pre-incubated with the mixtures indicated by the legend. The VHH-72-Fc used in this assay was purified after expression in ExpiCHO cells (n = 4). VHH-23-Fc is an irrelevant control VHH-Fc (n = 3). NI cells were not infected. Luciferase activity is reported in counts per second (c.p.s.) ± SEM.

## DISCUSSION

Here we report the isolation and characterization of two potently neutralizing single-domain antibodies from a llama immunized with prefusion-stabilized MERS-CoV and SARS-CoV-1 spikes. These VHHs bind to the spike RBDs with high affinity and are capable of neutralizing S pseudotyped viruses *in vitro*. To our knowledge, the isolation and characterization of SARS-CoV-1 S-directed VHHs have not been described before. Several MERS-CoV S-specific VHHs have been described, all of which have been directed against the RBD. Several of these VHHs have also been reported to block DPP4 binding, much like MERS VHH-55 (Stalin Raj et al., 2018; Zhao et al., 2018). By solving the crystal structures of these newly isolated VHHs in complex with their respective viral targets, we provide detailed insights into epitope binding and their mechanisms of neutralization.

A number of RBD-directed conventional antibodies have been described that are capable of neutralizing SARS-CoV-1 or MERS-CoV. The epitope of MERS VHH-55 overlaps with the epitopes of several of these MERS-CoV RBD-directed antibodies including C2, MCA1, m336, JC57-14, D12, 4C2 and MERS-27 (Chen et al., 2017; Li et al., 2015; Wang et al., 2018; Wang et al., 2015; Ying et al., 2015; Yu et al., 2015) (**S. Figure 7A**). The epitope of SARS VHH-72 does not significantly overlap with the epitopes of any previously described antibodies other than that of the recently described CR3022, which is also capable of binding to the RBDs of both SARS-CoV-1 and SARS-CoV-2 S (Hwang et al., 2006; Pak et al., 2009; Prabakaran et al., 2006; Walls et al., 2019; Yuan M, 2020) (**S. Figure 7B**). However, unlike SARS VHH-72, CR3022 does not prevent the binding of ACE2 and it lacks neutralizing activity against SARS-CoV-2 (Tian et al., 2020; Yuan M, 2020). Because SARS VHH-72 binds with nanomolar affinity to a portion of the SARS-CoV-1 S RBD that exhibits low sequence variation, as demonstrated by its cross-reactivity toward the WIV1-CoV and SARS-CoV-2 RBDs, it may broadly bind all S proteins from SARS-CoV-like viruses. We show that by engineering a bivalent VHH-72-Fc construct, we are able to compensate for the relatively high off-rate constant of the monovalent SARS VHH-72. This bivalent molecule expresses well in transiently transfected ExpiCHO cells (∼300 mg/L) and is capable of potently neutralizing SARS-CoV-2 S pseudoviruses *in vitro*.

Due to the inherent thermostability and chemostability of VHHs, they have been investigated as potential therapeutics against a number of diseases. Several HIV- and influenza-directed VHHs have been reported previously, and there are multiple RSV-directed VHHs that have been evaluated (Detalle et al., 2016; Ibanez et al., 2011; Koch et al., 2017; Rossey et al., 2017). The possibility of administering these molecules via a nebulized spray is particularly attractive in the case of respiratory pathogens because the VHHs could theoretically be inhaled directly to the site of infection in an effort to maximize bioavailability and function (Larios Mora et al., 2018). Due to the current lack of treatments for MERS, SARS and COVID-19 and the devastating effects associated with pandemic coronavirus outbreaks, both prophylactic and therapeutic interventions are sorely needed. It is our hope that due to their favorable biophysical properties and their potent neutralization capacity, MERS VHH-55, SARS VHH-72 and VHH-72-Fc may serve as both useful reagents for researchers and as potential therapeutic candidates.

## ACKNOWLEDGEMENTS

We thank members of the McLellan Laboratory for providing helpful comments on the manuscript. We would like to thank Dr. John Ludes-Meyers for assistance with cell transfection and protein production. This work was supported by a National Institutes of Health (NIH)/National Institute of Allergy and Infectious Disease (NIAID) grant R01-AI127521 (to J.S.M.). Research was supported by funding from VIB, Ghent University GOA project to N.C. and X.S., FWO and VLAIO fellowships and research projects to various VIB-CMB COVID-19 response team members. We acknowledge the team of the VIB Nanobody Service Facility for their services. D.D.V. was supported by a FWOsb fellowship, W.V. by the FWO-SBO grant “GlycoDelete”, S.P. by BMBF (Rapid consortium, 01K11723D), and B.S. by FWO-EOS project VIREOS. We are deeply indebted to the VIB-CMB COVID-19 response team members, who volunteered to offer their expertise and agile work under conditions of almost complete lockdown and societal standstill. We thank the support staff of both VIB-IRC and VIB-CMB centers, the members of VIB Discovery Sciences units’ COVID-19 team for rapid and consistent support and input. Argonne is operated by UChicago Argonne, LLC, for the US Department of Energy (DOE), Office of Biological and Environmental Research under Contract DE-AC02-06CH11357.

## AUTHOR CONTRIBUTIONS

Conceptualization, D.W., D.D.V., B.S.G., B.S., N.C., X.S., and J.S.M.; Investigation and visualization, D.W., D.D.V., K.S.C., G.M.T., W.V.B., K.R., L.v.S., M.H., S.P., and B.S.; Writing - Original Draft, D.W. and D.D.V.; Writing – Reviewing & Editing, D.W., D.D.V., K.S.C., G.M.T., B.S.G., N.C., B.S., X.S., and J.S.M.; Supervision, B.S.G., N.C., B.S., X.S., and J.S.M.

## MATERIALS AND METHODS

### Llama immunization

A llama, negative for antibodies against MERS-CoV and SARS-CoV-1 S glycoprotein, was subcutaneously immunized with approximately 150 µg recombinant SARS-CoV-1 S-2P protein on days 0, 7, 28 and 150 µg recombinant MERS-CoV S-2P protein on days 14 and 28 and 150 µg of both MERS-CoV S-2P and SARS-CoV-1 S-2P protein on day 35 (Kirchdoerfer et al., 2018; Pallesen et al., 2017). The adjuvant used was Gerbu LQ#3000. Immunizations and handling of the llama were performed according to directive 2010/63/EU of the European parliament for the protection of animals used for scientific purposes and approved by the Ethical Committee for Animal Experiments of the Vrije Universiteit Brussel (permit No. 13-601-1). Blood was collected 5 days after the last immunization for the preparation of lymphocytes. Total RNA from the peripheral blood lymphocytes was extracted and used as template for the first strand cDNA synthesis with oligo dT primer. Using this cDNA, the VHH encoding sequences were amplified by PCR and cloned between the *PstI* and *NotI* sites of the phagemid vector pMECS. In the pMECS vector, the VHH encoding sequence is followed by a linker, HA and His_6_ tag (AAAYPYDVPDYGSHHHHHH). Electro-competent E.coli TG1 cells were transformed with the recombinant pMECS vector resulting in a VHH library of about 3 x 10^8^ independent transformants. The resulting TG1 library stock was then infected with VCS M13 helper phages to obtain a library of VHH-presenting phages.

### Isolation of MERS- and SARS-CoV-directed VHH-displaying phages

Phages displaying MERS-CoV-specific VHHs were enriched after 2 rounds of biopanning on 20 μg of immobilized MERS-CoV S-2P protein in one well of a microtiter plate (type II, F96 Maxisorp, Nuc). For each panning round an uncoated well was used as a negative control. The wells were then washed 5 times with phosphate-buffered saline (PBS) + 0.05% Tween 20 and blocked with SEA BLOCK blocking buffer (Thermo Scientific) in the first panning round and 5% milk powder in PBS in the second panning round. About 10^11^ phages were added to the coated well and incubated for 1 hour at room temperature. Non-specifically bound phages were removed by washing with PBS + 0.05% Tween 20 (10 times in the first panning round and 15 times in the second panning round). The retained phages were eluted with TEA-solution (14% trimethylamine (Sigma) pH 10) and subsequently neutralized with 1 M Tris-HCl pH 8. The collected phages were amplified in exponentially growing *E.coli* TG1 cells, infected with VCS M13 helper phages and subsequently purified using PEG 8,000/NaCl precipitation for the next round of selection. Enrichment after each panning round was determined by infecting TG1 cells with 10-fold serial dilutions of the collected phages after which the bacteria were plated on LB agar plates with 100 μg mL^−1^ ampicillin and 1% glucose.

Phages displaying SARS-CoV-1 directed VHHs were enriched after 2 rounds of biopanning on 20 μg of SARS-CoV-1 S-2P protein captured with an anti-foldon antibody (generously provided by Dr. Vicente Mas) in one well of a microtiter plate (type II, F96 Maxisorp, Nuc). Before panning phages were first added to DS-Cav1 protein (McLellan et al., 2013) containing a C-terminal foldon domain, to deplete foldon specific phages. The unbound phages were next added to the coated well. Panning was performed as described above.

### Periplasmic ELISA screen to select MERS- and SARS-CoV directed VHHs

After panning, 45 individual colonies of phage infected bacteria isolated after the first panning round on MERS-CoV S-2P or SARS-CoV-1 S-2P protein and 45 individual colonies isolated after the second panning round on MERS-CoV S-2P or SARS-CoV-1 S-2P protein were randomly selected for further analysis by ELISA for the presence of MERS-CoV and SARS-CoV-1 specific VHHs, respectively. The individual colonies were inoculated in 2 mL of terrific broth (TB) medium with 100 µg/mL ampicillin in 24-well deep well plates. After growing individual colonies for 5 hours at 37 °C, isopropyl β-D-1-thiogalactopyranoside (IPTG) (1 mM) was added to induce VHH expression during overnight incubation at 37 °C. To prepare periplasmic extract, the bacterial cells were pelleted and resuspended in 250 μL TES buffer (0.2 M Tris-HCl pH 8, 0.5 mM EDTA, 0.5 M sucrose) and incubated at 4 °C for 30 min. Subsequently 350 μL water was added to induce an osmotic shock. After 1 hour incubation at 4 °C followed by centrifugation, the periplasmic extract was collected.

VHH-containing periplasmic extracts were then tested for binding to either MERS-CoV S-2P or SARS-CoV-1 S-2P protein. Briefly, in the PE-ELISA screen after panning on MERS-CoV S-2P protein, wells of microtiter plates (type II, F96 Maxisorp, Nuc) were coated overnight at 37°C with 100 ng MERS-CoV S-2P (without foldon), MERS-CoV S-2P protein (with foldon) or as negative controls coated with SARS-CoV-1 S-2P protein (with foldon), HCoV-HKU1 S-2P (without foldon), DS-Cav1 (with foldon) or bovine serum albumin (BSA, Sigma-Aldrich). In the PE-ELISA screen after panning on SARS-CoV-1 S protein wells of microtiter plates (type II, F96 Maxisorp, Nuc) were coated with 100 ng SARS-CoV-1 S-2P protein (with foldon), SARS-CoV-1 S-2P protein captured with an anti-foldon antibody (with foldon) or as negative controls coated with MERS-CoV S-2P (without foldon), HCoV-HKU1 S-2P (without foldon), DS-Cav1 (with foldon) or bovine serum albumin (BSA, Sigma-Aldrich). The coated plates were blocked with 5% milk powder in PBS and 50 μL of the periplasmic extract was added to the wells. Bound VHHs were detected with anti-HA (1/2,000, MMS-101P Biolegend) mAb followed by horseradish peroxidase (HRP)-linked anti-mouse IgG (1/2,000, NXA931, GE Healthcare). Periplasmic fractions, for which the OD_450_ value of the antigen coated wells were at least two times higher than the OD_450_ value of the BSA coated wells, were considered to be specific for the coated antigen and selected for sequencing. The selected clones were grown in 3 mL of LB medium with 100 μg/mL ampicillin. The DNA of the selected colonies was isolated using the QIAprep Spin Miniprep kit (Qiagen) and sequenced using the MP057 primer (5’-TTATGCTTCCGGCTCGTATG-3’).

### Cloning of MERS- and SARS-CoV directed VHHs into a Pichia pastoris expression vector

In order to express the MERS- and SARS-CoV VHHs in *Pichia pastoris*, the VHH encoding sequences were cloned in the pKai61 expression vector (Schoonooghe et al., 2009). In the vector, the VHH sequences contain a C-terminal 6x His-tag, are under the control of the methanol inducible AOX1 promotor and in frame with a modified version of the *S.cerevisae* α-mating factor prepro signal sequence. The vector contains a Zeocine resistance marker for selection in bacteria as well as in yeast cells. The VHH encoding sequences were amplified by PCR using the following forward and reverse primer (5’-GGCGGGTATCTCTCGAGAAAAGGCAGGTGCAGCTGCAGGAGTCTGGG-3’) and (5’-CTAACTAGTCTAGTGATGGTGATGGTGGTGGCTGGAGACGGTGACCTGG-3’) and cloned between the *Xho*I and *Spe*I sites in the pKai61 vector. The vectors were linearized by *PmeI* and transformed in the *Pichia pastoris* strain GS115 using the condensed transformation protocol described by Lin-Cereghino *et al* (Lin-Cereghino et al., 2005). After transformation, the yeast cells were plated on YPD plates (1% (w/v) yeast extract, 2% (w/v) peptone, 2% (w/v) dextrose and 2% (w/v) agar) supplemented with zeocin (100 µg/mL) for selection.

### Generation of bivalent VHH-constructs for production in *Pichia pastoris*

To generate bivalent tandem tail-to-head VHH constructs, the VHH sequence was amplified by PCR using the following forward (5’-GGGGTATCTCTCGAGAAAAGGCAGGTGCAGCTGGTGGAGTCTGGG-3’) and reverse (5’-AGACTCCTGCAGCTGCACCTGACTACCGCCGCCTCCAGATCCACCTCCGCCACTACCGCCTCCGCCGCTGGAGACGGTGAC CTGGG-3’) primers, thereby removing a *Pst*I site from the beginning of the VHH coding sequence and adding a (GGGGS)_3_ linker and the start of the VHH coding sequence with a *Pst*I site at the end of the sequence. After PCR, the fragment was cloned between the *XhoI* and *SpeI* sites in a SARS VHH-72 containing pKai61 vector, thereby generating a homo-bivalent construct. The vector containing this bivalent VHH was linearized and transformed in GS155 *Pichia pastoris* cells as outlined above.

### Pichia production and purification MERS- and SARS-CoV directed VHHs

The transformed *Pichia pastoris* clones were first expressed in 2 mL cultures. On day 1, 4 clones of each construct were inoculated in 2 mL of YPNG medium (2% pepton, 1% Bacto yeast extract, 1.34% YNB, 0.1 M potassium phosphate pH 6, 0.00004% biotin, 1% glycerol) with 100 μg/mL Zeocin (Life Technologies) and incubated while shaking at 28 °C for 24 hours. The next day, the cells were pelleted by centrifugation and the medium was replaced by YPNM medium (2% pepton, 1% Bacto yeast extract, 1.34% YNB, 0.1 M potassium phosphate pH 6.0, 1% methanol) to induce VHH expression. Cultures were incubated at 28 °C and 50 μL of 50% methanol was added at 16, 24 and 40 h. After 48 h, the yeast cells were pelleted and the supernatant was collected. The presence of soluble VHHs in the supernatants was verified using SDS-PAGE and subsequent Coomassie Blue staining. VHH-containing supernatants of the different clones for each construct were pooled and the VHHs were purified using HisPur^TM^ Ni-NTA Spin Plates (88230, Thermo Scientific™). Next, purified VHHs were concentrated on AcroPrep^TM^ Advance 96-well filter plates for ultrafiltration 3 kDa cutoff (8033,Pall) and the imidazole-containing elution buffer was exchanged with PBS.

Production was scaled up (50 mL) for the VHHs with neutralizing capacity. Growth and methanol induction conditions and harvesting of medium were similar as mentioned above for the 2 mL cultures. The secreted VHHs in the medium were precipitated by ammonium sulfate (NH_4_)_2_SO_4_ precipitation (80% saturation) for 4 h at 4 °C. The insoluble fraction was pelleted by centrifugation at 20,000 g and resuspended in 10 mL binding buffer (20 mM NaH_2_PO_4_ pH 7.5, 0.5M NaCl and 20 mM imidazole pH 7.4). The VHHs were purified from the solution using a 1 mL HisTrap HP column (GE Healthcare). To elute the bound VHHs a linear imidazole gradient starting from 20 mM and ending at 500 mM imidazole in binding buffer over a total volume of 20 mL was used. VHH containing fractions were pooled and concentrated and the elution buffer was exchanged with PBS with a Vivaspin column (5 kDa cutoff, GE Healthcare).

### Enzyme-linked immunosorbent assay

Wells of microtiter plates (type II, F96 Maxisorp, Nuc) were coated overnight at 4 °C, respectively, with 100 ng recombinant MERS-CoV S-2P protein (with foldon), SARS-CoV-1 S-2P protein (with foldon), MERS-CoV RBD, MERS-CoV NTD, MERS-CoV S1, SARS-CoV-1 RBD, SARS-CoV-1 NTD or Fc-tagged SARS-CoV-2 RBD-SD1. The coated plates were blocked with 5% milk powder in PBS. Dilution series of the VHHs were added to the wells. Binding was detected by incubating the plates sequentially with either mouse anti-Histidine Tag antibody (MCA1396, Abd Serotec) followed horseradish peroxidase (HRP)-linked anti-mouse IgG (1/2000, NXA931, GE Healthcare) or Streptavidin-HRP (554066, BD Biosciences) or by an HRP-linked rabbit anti-camelid VHH monoclonal antibody (A01861-200, GenScript). After washing 50 µL of TMB substrate (Tetramethylbenzidine, BD OptETA) was added to the plates and the reaction was stopped by addition of 50 µL of 1 M H_2_SO4. The absorbance at 450 nM was measured with an iMark Microplate Absorbance Reader (Bio Rad). Curve fitting was performed using nonlinear regression (Graphpad 7.0).

### CoV pseudovirus neutralization

Pseudovirus neutralization assay methods have been previously described (Pallesen et al., 2017; Wang et al., 2015). Briefly, pseudoviruses expressing spike genes for MERS-CoV England1 (GenBank ID: AFY13307) and SARS-CoV-1 Urbani (GenBank ID: AAP13441.1) were produced by co-transfection of plasmids encoding a luciferase reporter, lentivirus backbone, and spike genes in 293T cells (Wang et al., 2015). Serial dilutions of VHHs were mixed with pseudoviruses, incubated for 30 min at room temperature, and then added to previously-plated Huh7.5 cells. 72 hours later, cells were lysed, and relative luciferase activity was measured. Percent neutralization was calculated considering uninfected cells as 100% neutralization and cells transduced with only pseudovirus as 0% neutralization. IC_50_ titers were determined based on sigmoidal nonlinear regression.

Replication-deficient VSV pseudotyped with MERS-CoV S, SARS-CoV-1 S or SARS-CoV-2 S and coding for GFP or firefly luciferase were generated as described previously (Berger Rentsch and Zimmer, 2011; Hoffmann, 2020). For the VSV pseudotype neutralization experiments, the pseudoviruses were incubated for 30 min at 37 °C with different dilutions of purified VHHs or with dilution series of culture supernatant of 293S cells that had been transfected with plasmids coding for SARS VHH-72 fused to human IgG1 Fc (VHH-72-Fc) or with GFP-binding protein (GBP: a VHH specific for GFP). The incubated pseudoviruses were subsequently added to confluent monolayers of Vero E6 cells. Sixteen hours later, the transduction efficiency was quantified by measuring the firefly luciferase activity in cell lysates using the firefly luciferase substrate of the dual-luciferase reporter assay system (Promega) and a Glowmax plate luminometer (Promega).

### Mammalian protein expression and purification

Mammalian expression plasmids encoding for SARS VHH72, MERS VHH55, residues 367-589 of MERS-CoV S (England1 strain), residues 320-502 of SARS-CoV-1 S (Tor2 strain), residues 307-510 of WIV1-CoV S, residues 319-591 of SARS-CoV-2 S, residues 1-281 of SARS-CoV-1 S (Tor2 strain), residues 1-351 of MERS-CoV S (England1 strain), residues 1-751 of MERS-CoV S (England1 strain), residues 1-1190 of SARS-CoV-1 S (Tor2 strain) with K968P and V969P substitutions (SARS-CoV-1 S-2P), residues 1-1291 of MERS-CoV S (England1 strain) with V1060P and L1061P substitutions (MERS S-2P), residues 1-1208 of SARS-CoV-2 S with K986P and V987P substitutions (SARS-CoV-2 S-2P), residues 1-615 of ACE2 and residues 40-766 of DPP4 were transfected into FreeStyle293 cells using polyethylenimine (PEI). All of these plasmids contained N-terminal signal sequences to ensure secretion into the cell supernatant.

Supernatants were harvested and constructs containing C-terminal HRV3C cleavage sites, 8x His-Tags and Twin-Strep-Tags (SARS VHH72, MERS VHH55, MERS-CoV S1, SARS-CoV-1 S-2P, MERS-CoV S-2P, SARS-CoV-2 S-2P, ACE2 and DPP4) were purified using Strep-Tactin resin (IBA). Constructs containing C-terminal HRV3C cleavage sites and Fc-tags (SARS-CoV-1 RBD, MERS-CoV RBD, WIV1-CoV RBD, SARS-CoV-2 RBD-SD1, SARS-CoV-1 NTD, MERS-CoV NTD) were purified using Protein A resin (Pierce). The SARS-CoV-1 RBD, MERS-CoV RBD, WIV1-CoV RBD, SARS-CoV-2 RBD-SD1, SARS VHH-72, MERS VHH-55, MERS-CoV NTD and SARS-CoV-1 NTD were then further purified using a Superdex 75 column (GE Healthcare) in 2 mM Tris pH 8.0, 200 mM NaCl and 0.02% NaN_3_. MERS-CoV S1, SARS-CoV-1 S-2P, MERS-CoV S-2P, ACE2 and DPP4 were further purified using a Superose 6 column (GE Healthcare) in 2 mM Tris pH 8.0, 200 mM NaCl and 0.02% NaN_3_.

HEK 293S cells were transfected with VHH-72-Fc or VHH-72-Fc (S) encoding plasmids using PEI. Briefly, suspension-adapted and serum-free HEK 293S cells were seeded at 3 x 10^6^ cells/mL in Freestyle-293 medium (ThermoFisher Scientific). Next, 4.5 µg of pcDNA3.3-VHH72-Fc plasmid DNA was added to the cells and incubated on a shaking platform at 37 °C and 8% CO_2_, for 5 min. Next, 9 µg of PEI was added to the cultures, and cells were further incubated for 5 h, after which an equal culture volume of Ex-Cell-293 (Sigma) was added to the cells. Transfections were incubated for 4 days, after which cells were pelleted (10’, 300g) and supernatants were filtered before further use.

VHH-72-Fc was expressed in ExpiCHO cells (ThermoFisher Scientific), according to the manufacturer’s protocol. Briefly, a 25 mL culture of 6 x10^6^ cells/mL, grown at 37 °C and 8% CO_2_ was transfected with 20 µg of pcDNA3.3-VHH-72-Fc plasmid DNA using ExpiFectamine CHO reagent. One day after transfection, 150 µL of ExpiCHO enhancer and 4 mL of ExpiCHO feed was added to the cells, and cultures were further incubated at 32 °C and 5% CO_2_. Cells were fed a second time 5 days post-transfection. Cultures were harvested as soon as cell viability dropped below 75%. For purification of the VHH-72-Fc, supernatants were loaded on a 5 mL MabSelect SuRe column (GE Healthcare). Unbound proteins were washed away with McIlvaine buffer pH 7.2, and bound proteins were eluted using McIlvaine buffer pH 3. Immediately after elution, protein-containing fractions were neutralized using a saturated Na_3_PO_4_ buffer. These neutralized fractions were then pooled, and loaded onto a HiPrep Desalting column for buffer exchange into storage buffer (25 mM L-Histidine, 125 mM NaCl).

### Surface plasmon resonance

His-tagged SARS VHH-72 or MERS VHH-55 was immobilized to a single flow cell of an NTA sensorchip at a level of ∼400 response units (RUs) per cycle using a Biacore X100 (GE Healthcare). The chip was doubly regenerated using 0.35 M EDTA and 0.1 M NaOH followed by 0.5 mM NiCl_2_. Three samples containing only running buffer, composed of 10 mM HEPES pH 8.0, 150 mM NaCl and 0.005% Tween 20, were injected over both ligand and reference flow cells, followed by either SARS-CoV-1 RBD, WIV1-CoV RBD, SARS-CoV-2 RBD-SD1 or MERS-CoV RBD serially diluted from 50-1.56 nM, with a replicate of the 3.1 nM concentration. The resulting data were double-reference subtracted and fit to a 1:1 binding model using the Biacore X100 Evaluation software.

### Crystallization and data collection

Plasmids encoding for MERS VHH-55 and residues 367-589 of MERS-CoV S with a C-terminal HRV3C cleavage site and a monomeric human Fc tag were co-transfected into kifunensin-treated FreeStyle 293F cells, as described above. After purifying the cell supernatant with Protein A resin, the immobilized complex was treated with HRV3C protease and Endoglycosidase H to remove both tags and glycans. The complex was then purified using a Superdex 75 column in 2 mM Tris pH 8.0, 200 mM NaCl and 0.02% NaN_3_. The purified complex was then concentrated to 5.0 mg/mL and used to prepare hanging-drop crystallization trays. Crystals grown in 1.0 M Na/K phosphate pH 7.5 were soaked in mother liquor supplemented with 20% ethylene glycol and frozen in liquid nitrogen. Diffraction data were collected to a resolution of 3.40 Å at the SBC beamline 19-ID (APS, Argonne National Laboratory) Plasmids encoding for SARS VHH-72 and residues 320-502 of SARS-CoV-1 S with a C-terminal HRV3C cleavage site and a monomeric human Fc tag were co-transfected into kifunensin-treated FreeStyle 293F cells, as described above. After purifying the cell supernatant with Protein A resin, the immobilized complex was treated with HRV3C protease and Endoglycosidase H to remove both tags and glycans. The processed complex was subjected to size-exclusion chromatography using a Superdex 75 column in 2 mM Tris pH 8.0, 200 mM NaCl and 0.02% NaN_3_. The purified complex was then concentrated to 10.0 mg/mL and used to prepare hanging-drop crystallization trays. Crystals grown in 0.1 M Tris pH 8.5, 0.2 M LiSO_4_, 0.1 M LiCl and 8% PEG 8000 were soaked in mother liquor supplemented with 20% glycerol and frozen in liquid nitrogen. Diffraction data were collected to a resolution of 2.20 Å at the SBC beamline 19-ID (APS, Argonne National Laboratory)

### Structure determination

Diffraction data for both complexes were indexed and integrated using iMOSFLM before being scaled in AIMLESS (Battye et al., 2011; Evans and Murshudov, 2013). The SARS-CoV-1 RBD+SARS VHH-72 dataset was phased by molecular replacement in PhaserMR using coordinates from PDBs 2AJF and 5F1O as search ensembles (McCoy, 2007). The MERS-CoV RBD+MERS VHH-55 dataset was also phased by molecular replacement in PhaserMR using coordinates from PDBs 4L72 and 5F1O as search ensembles. The resulting molecular replacement solutions were iteratively rebuilt and refined using Coot, ISOLDE and Phenix (Adams et al., 2002; Croll, 2018; Emsley and Cowtan, 2004). Crystallographic software packages were curated by SBGrid (Morin et al., 2013).

### Biolayer interferometry

Anti-human capture (AHC) tips (FortéBio) were soaked in running buffer composed of 10 mM HEPES pH 7.5, 150 mM NaCl, 3 mM EDTA, 0.005% Tween 20 and 1 mg/mL BSA for 20 min before being used to capture either Fc-tagged SARS-CoV-1 RBD, Fc-tagged SARS-CoV-2 RBD-SD1 or Fc-tagged MERS-CoV RBD to a level of 0.8 nm in an Octet RED96 (FortéBio). Tips were then dipped into either 100 nM MERS VHH-55 or 100 nM SARS VHH-72. Tips were next dipped into wells containing either 1 μM ACE2 or 100 nM DPP4 supplemented with the nanobody that the tip had already been dipped into to ensure continued saturation. Data were reference-subtracted and aligned to each other in Octet Data Analysis software v11.1 (FortéBio) based on a baseline measurement that was taken before being dipped into the final set of wells that contained either ACE2 or DPP4.

BLI measurements were also performed with VHH-72-Fc fusion produced in HEK 293S cells. SARS-CoV-2 RBD with a mouse IgG1 Fc tag (Sino Biological) was immobilized to an anti-mouse IgG Fc capture (AMC) tip (FortéBio) to a response level of 0.5 nm. Supernatant of non-transfected and VHH-72-Fc transfected HEK293-S cells was applied in a three-fold dilution series in kinetics buffer. Binding was measured at 30 °C, with baseline and dissociation measured in equal dilution of non-transformed HEK293S supernatant in kinetics buffer. Between analyses, biosensors were regenerated by three times 20 s exposure to regeneration buffer (10 mM glycine pH 1.7).

### Flow cytometry

Binding of VHH-72-Fc, VHH-72-Fc (S) and monomeric SARS VHH-72 to SARS-CoV-1 and SARS-CoV-2 S was analyzed by flow cytometry using cells transfected with a GFP expression plasmid combined with an expression plasmid for either SARS-CoV-1 or SARS-Cov-2 S. HEK 293S culture media (1/20 diluted in PBS + 0.5%BSA) of VHH-72-Fc and VHH-72-Fc (S) transformants were incubated with transfected cells. Binding of the VHH-72-Fc and VHH-72-Fc (S) to cells was determined by an AF633 conjugated goat anti-human IgG antibody and binding to SARS-CoV-1 or SARS-CoV-2 S was calculated as the mean AF633 fluorescence intensity (MFI) of GFP expressing cells (GFP^+^) divided by the MFI of GFP negative cells (GFP^-^).

### RBD competition assay on Vero E6 cells

SARS-CoV-2 RBD fused to murine IgG Fc (Sino Biological) at a final concentration of 0.4 μg/mL was incubated with a dilution series of tail-to-head bivalent VHHs or VHH-Fc fusions and incubated at room temperature for 20 min before an additional 10 min incubation on ice. Vero E6 cells grown at sub-confluency were detached by cell dissociation buffer (Sigma) and trypsin treatment. After washing once with PBS the cells were blocked with 1% BSA in PBS on ice. All remaining steps were also performed on ice. The mixtures containing RBD and tail-to-head bivalent VHHs or VHH-Fc fusions were added to the cells and incubated for one hour.

Subsequently, the cells were washed 3 times with PBS containing 0.5% BSA and stained with an AF647 conjugated donkey anti-mouse IgG antibody (Invitrogen) for 1 hour. Following additional 3 washes with PBS containing 0.5% BSA, the cells were analyzed by flow cytometry using an BD LSRII flow cytometer (BD Biosciences).

---

**Supplementary Figure 1:**
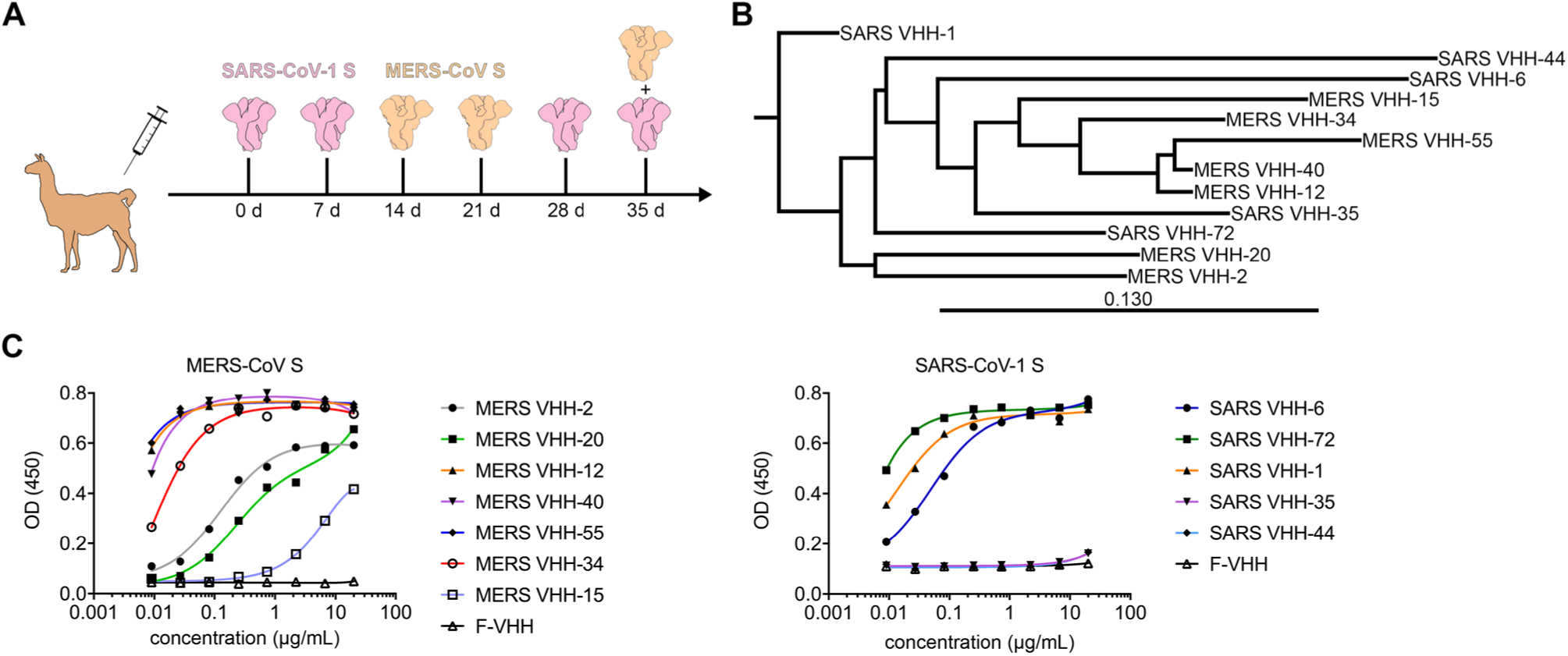
CoV VHH immunization and panning. **A**) Schematic depicting the immunization strategy that was used to isolate both SARS-CoV-1 S and MERS-CoV S-directed VHHs from a single llama. The prefusion stabilized SARS-CoV-1 spike is shown in pink and the prefusion stabilized MERS-CoV spike is shown in tan. **B**) Phylogenetic tree of the isolated MERS-CoV and SARS-CoV S-directed VHHs, based on the neighbor joining method. **C**) Reactivity of MERS-CoV and SARS-CoV S-directed VHHs against the prefusion stabilized MERS-CoV S and SARS-CoV-1 S protein, respectively. A VHH against an irrelevant antigen (F-VHH) was included as a control.

**Supplementary Figure 2:**
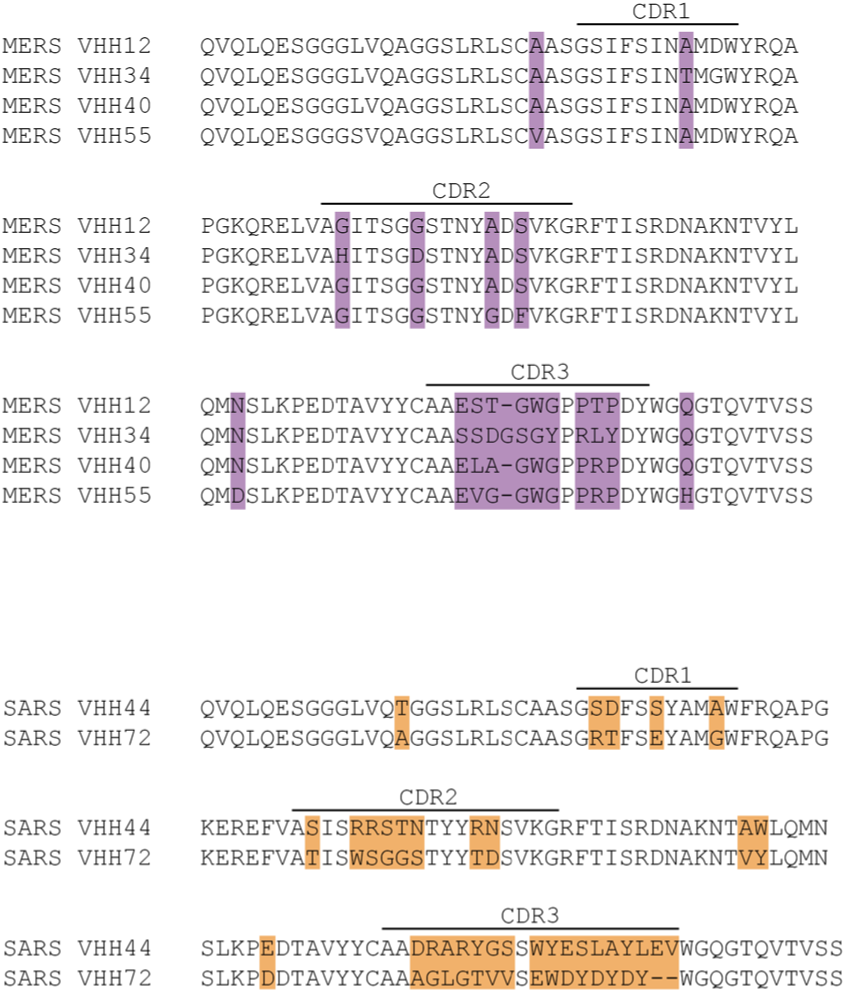
Sequence alignment of neutralizing SARS-CoV and MERS-CoV S-directed VHHs. Invariant residues are shown as black dots. The CDRs are shown underneath black lines..

**Supplementary Figure 3:**
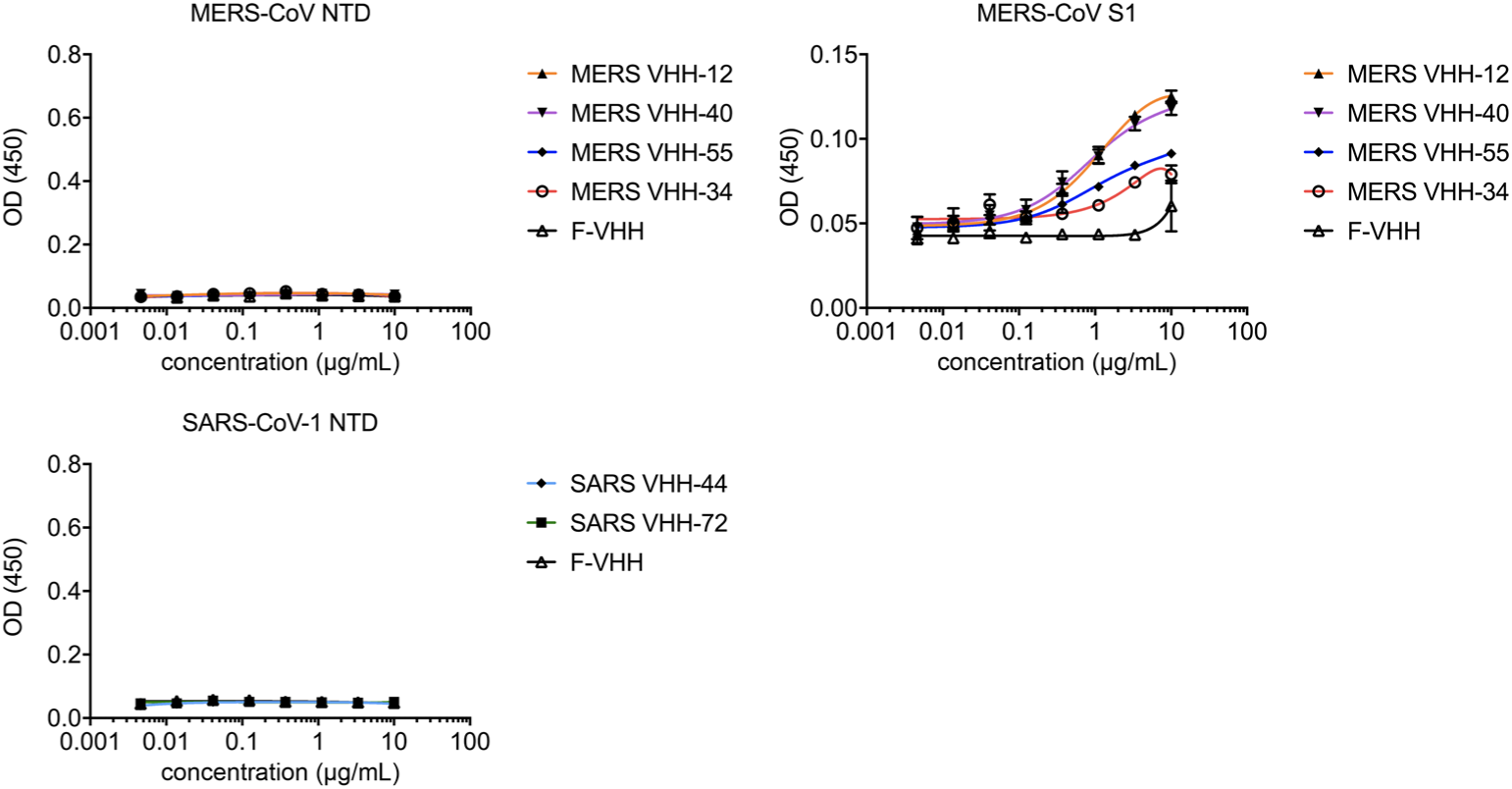
Lack of binding of MERS-CoV and SARS-CoV directed VHHs to non-RBD epitopes. ELISA data showing binding of the MERS-CoV specific VHHs to the MERS-CoV S1 protein and absence of binding of the MERS-CoV and SARS-CoV specific VHHs against the MERS-CoV NTD and SARS-CoV-1 NTD, respectively. A VHH against an irrelevant antigen (F-VHH) was included as a control.

**Supplementary Figure 4:**
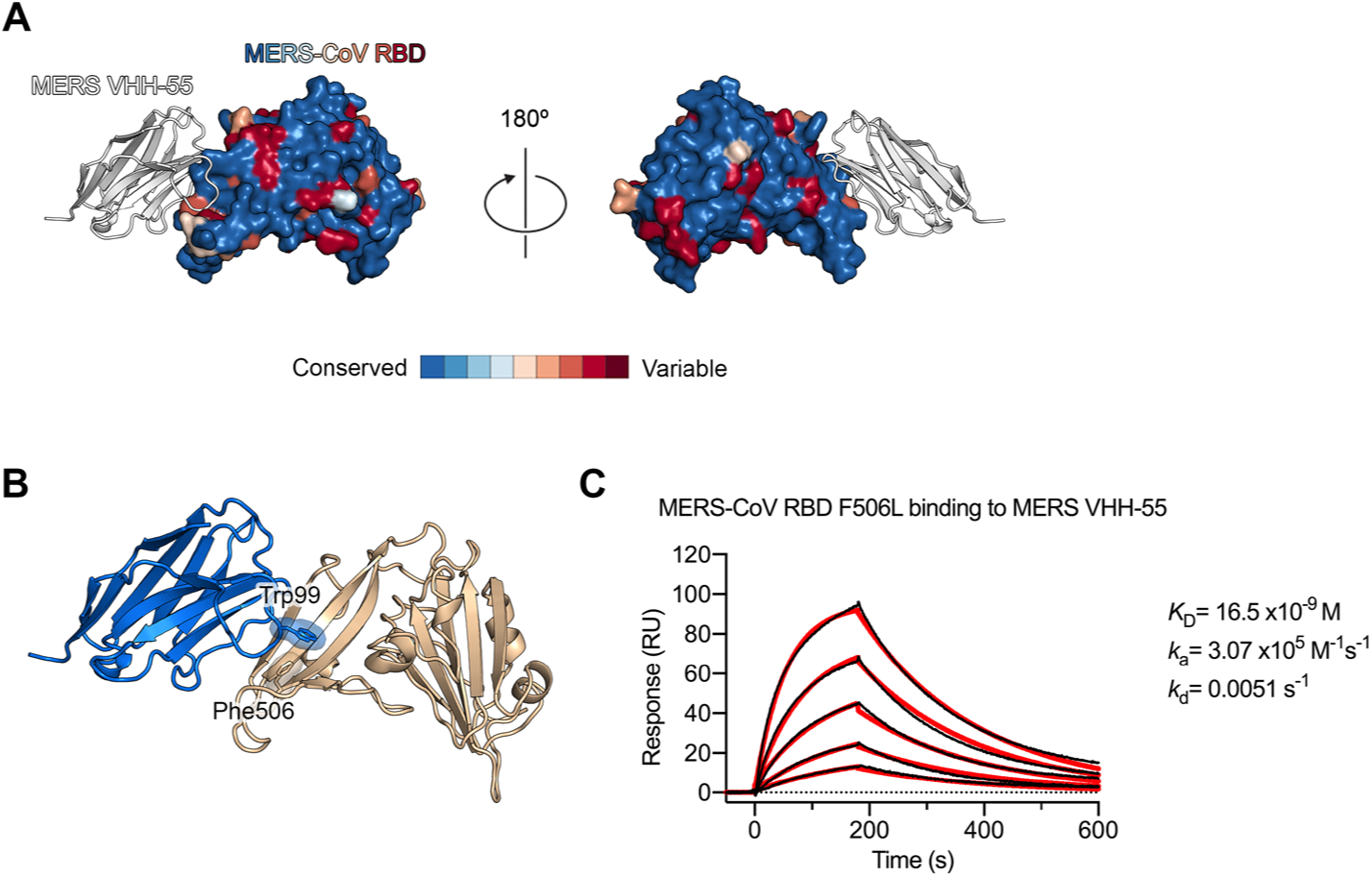
MERS VHH-55 binds to a relatively conserved epitope on the MERS-CoV RBD. A) The crystal structure of MERS VHH-55 bound to the MERS-CoV RBD is shown with MERS VHH-55 in white ribbons and the MERS-CoV RBD as a multicolored molecular surface. More variable residues are shown in warm colors and more conserved residues are shown in cool colors according to the spectrum (*bottom*). Sequence alignments and variability mapping was performed using ConSurf. **B**) The crystal structure of MERS VHH-55 bound to the MERS-CoV RBD is shown as ribbons with MERS VHH-55 colored blue and the MERS-CoV RBD colored tan. Phe506 from the MERS-CoV RBD and Trp99 from MERS VHH-55, which are thought to form hydrophobic interactions with one another are shown as sticks surrounded by a transparent molecular surface. **C**) SPR sensorgram measuring the binding of MERS VHH-55 to the naturally occurring MERS-CoV RBD F506L variant. Binding curves are colored black and the fit of the data to a 1:1 binding model is colored red.

**Supplementary Figure 5:**
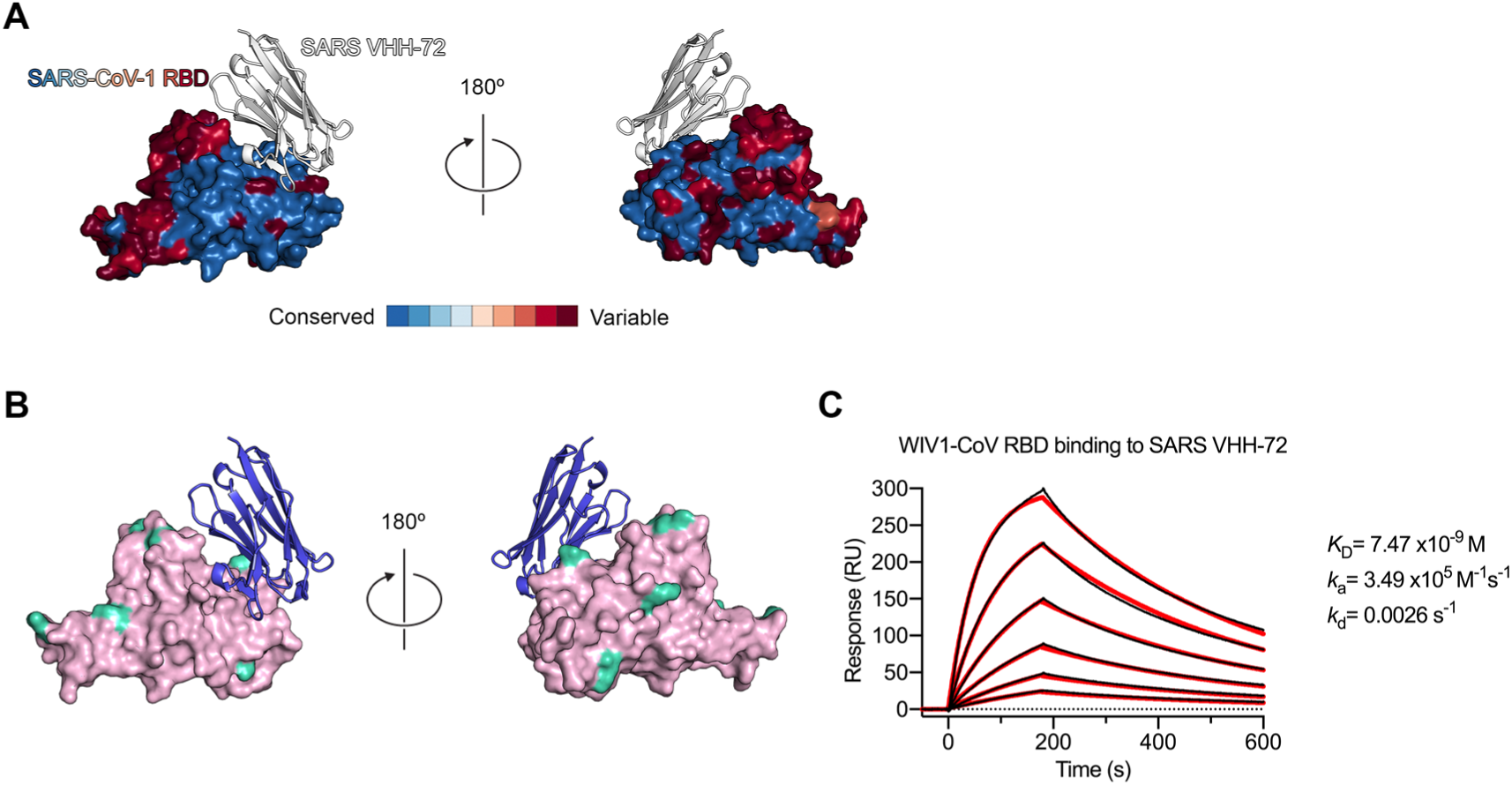
SARS VHH-72 binds to a broadly conserved epitope on the SARS-CoV-1 RBD. **A**) The crystal structure of SARS VHH-72 bound to the SARS-CoV-1 RBD is shown, with colors corresponding to those of **S**Fig 4A. **B**) The crystal structure of SARS VHH-72 bound to the SARS-CoV-1 RBD is shown with SARS VHH-72 as dark blue ribbons and the RBD as a pink molecular surface. Amino acids that vary between SARS-CoV-1 and WIV1-CoV are colored teal. **C**) SPR sensorgram measuring the binding of SARS VHH-72 to the WIV1-CoV RBD. Binding curves are colored black and the fit of the data to a 1:1 binding model is colored red.

**Supplementary Figure 6:**
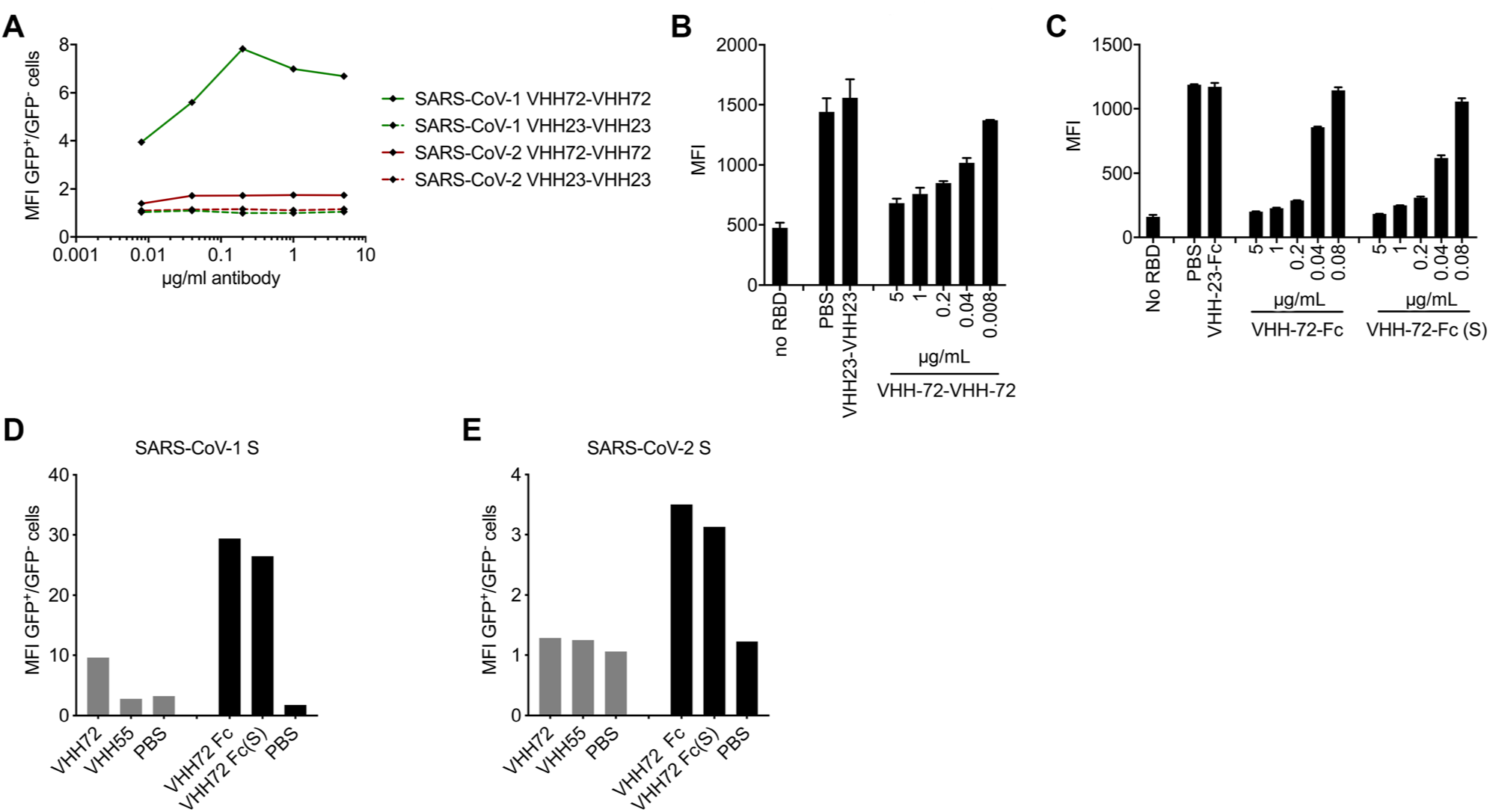
Engineering a functional bivalent VHH construct. **A**) Flow cytometry measuring the binding of the bivalent SARS VHH-72 tail-to-head fusion (VHH-72-VHH-72) to SARS-CoV-1 or SARS-CoV-2 S expressed on the cell surface. VHH-23-VHH-23, a bivalent tail-to-head fusion of an irrelevant nanobody, was included as a negative control. **B**) Binding of SARS-CoV-2 RBD-SD1 to Vero E6 cells is prevented by VHH-72-VHH-72 in a dose-dependent fashion. Binding of SARS-CoV-2 RBD-SD1 to Vero E6 cells was detected by flow cytometry in the presence of the indicated bivalent VHHs (n = 2 except VHH-72-VHH-72 and VHH-23-VHH-23 at 5 µg/ml, n = 5). **C**) Binding of SARS-CoV-2 RBD-SD1 to Vero E6 cells is prevented by bivalent VHH-72-Fc fusion proteins in a dose-dependent fashion. Binding of SARS-CoV-2 RBD-SD1-Fc to Vero E6 cells was detected by flow cytometry in the presence of the indicated constructs and amounts (n = 2 except no RBD, n = 4). **D**) Cell surface binding of SARS VHH-72 to SARS-CoV-1 S. 293T cells were transfected with a GFP expression plasmid together with a SARS-CoV-1 S expression plasmid. Binding of the indicated protein is expressed as the median fluorescent intensity (MFI), measured to detect the His-tagged MERS VHH-55 or SARS VHH-72 or the SARS VHH-72-Fc fusions, of the GFP positive cells divided by the MFI of the GFP negative cells**. E)** Cell surface binding of SARS VHH-72 to SARS-CoV-2. MFI was calculated using the same equation as **S.** Figure 6D.

**Supplementary Figure 7:**
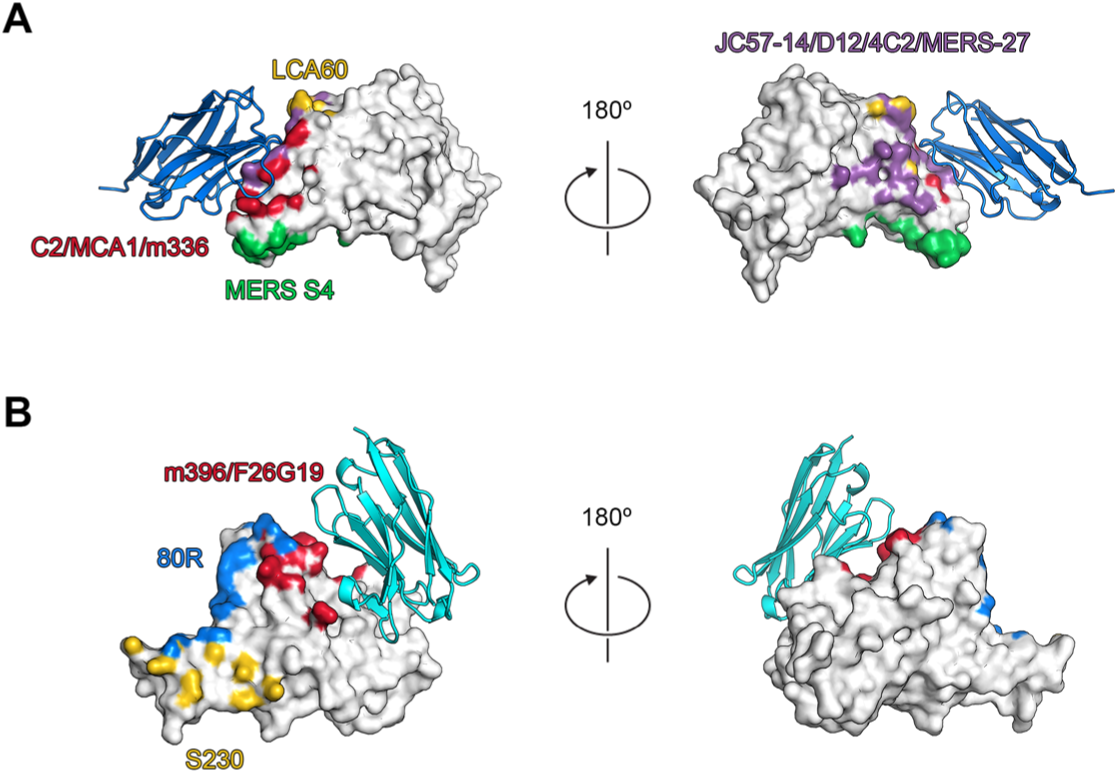
Comparison of the CoV VHH epitopes with known RBD-directed antibodies. **A**) The structure of MERS VHH-55 bound to the MERS-CoV RBD is shown with MERS VHH-55 as blue ribbons and the MERS-CoV RBD as a white molecular surface. Epitopes from previously reported crystal structures of the MERS-CoV RBD bound by RBD-directed antibodies are shown as colored patches on the MERS-CoV RBD surface. The LCA60 epitope is shown in yellow, the MERS S4 epitope is shown in green, the overlapping C2/MCA1/m336 epitopes are shown in red and the overlapping JC57-14/D12/4C2/MERS-27 epitopes are shown in purple. **B**) The structure of SARS VHH-72 bound to the SARS-CoV-1 RBD is shown with SARS VHH-72 as cyan ribbons and the SARS-CoV-1 RBD as a white molecular surface. Epitopes from previously reported crystal structures of the SARS-CoV-1 RBD bound by RBD-directed antibodies are shown as colored patches on the SARS-CoV-1 RBD surface. The 80R epitope is shown in blue, the S230 epitope is shown in yellow, and the overlapping m396/F26G19 epitopes are shown in red.

